# Functional Genomics in Human iPSC-Derived Sinoatrial Node Cells Reveals Shared Regulatory Determinants of Heart Rate and Atrial Fibrillation

**DOI:** 10.1101/2023.07.01.547335

**Authors:** James L. Engel, Xianglong Zhang, Rajiv A. Mohan, Navraj Lally, Amanda W. Soe, Grace Yang, Daniel R. Lu, Olaia F. Vila, Vanessa Arias, Jasper Lee, Christophero Hale, Yi-Hsiang Hsu, Chi-Ming Li, Gabriel B. Loeb, Arun Padmanabhan, Roland S. Wu, Yen-Sin Ang, Vasanth Vedantham

**Author notes:** <u>Corresponding Authors:</u> Yen-Sin Ang, PhD, Vasanth Vedantham, MD, PhD, Smith Cardiovascular Research Building 555 Mission Bay Blvd South, 352M, San Francisco, CA 94158. These authors contributed equally (James L. Engel, Xianglong Zhang).

## Abstract

Human model systems for functional genomics of heart rhythm are needed to translate genome wide association studies into biological insight and actionable targets. Here we develop a human induced pluripotent stem cell sinoatrial node system that recapitulated the transcriptional and epigenetic heterogeneity of primary human pacemaker tissue, permitting exploration of heart rhythm-associated single nucleotide polymorphisms (SNPs) in a cell subtype-specific manner. Using self-transcribing active regulatory region sequencing (STARR-seq), we experimentally validated numerous enhancers containing heart rhythm associated variants. We demonstrated the utility of this platform for fine mapping of candidate causal SNPs by identifying an AF-associated variant at the *ATXN1* locus that affects signal responsiveness of an enhancer, and a variant at the *GNB4* locus that regulates cardiac autonomic sensitivity, leading to a pleiotropic effect on heart rate and atrial fibrillation. Taken together, these data establish a robust human cellular system to explore the mechanistic basis of heart rhythm heritability.

## INTRODUCTION

Pacemaker cardiomyocytes (PCs) in the sinoatrial node (SAN) generate a synchronized electrical wavefront that initiates the heartbeat^1^. PCs are integrated with the autonomic nervous system and have a complex functional and anatomical organization, allowing the SAN to achieve beat-to-beat modulation of heart rate to match changes in metabolic demand^2,3^. The SAN head, located at the junction of the right atrium and superior vena cava, is most sensitive to autonomic input, driving impulse generation under high sympathetic tone^4,5^. The more inferiorly located SAN tail stabilizes basal automaticity when the SAN head is inhibited by high parasympathetic tone, while a transition zone at the interface of the SAN and atrial myocardium promotes unidirectional impulse transmission from the SAN to the atrium.^2,6,7^

SAN dysfunction (SND), among the most common heart rhythm disorders, causes slow heartbeat leading to a need for pacemaker implantation. While it is well established that SAN head, SAN tail, and atrial-like transitional cells engage different transcriptional networks,^7–10^ the detailed mechanisms that drive cell subtype-specific gene programs underlying normal SAN function and SND are incompletely understood. Heart rate is a strongly heritable trait^11^, and genome wide association studies have defined hundreds of loci associated with heart rate determination and susceptibility to SND,^12,13^ providing a potential window into the cellular and molecular basis for SAN function and SND. However, the cellular subtypes and specific contexts in which common variants regulate heart rhythm and disease susceptibility are still not well-established, partly due to the lack of suitable experimental model systems.

Furthermore, SND and slow heart rate are associated with a higher incidence of atrial fibrillation (AF) ^14^, a distinct arrhythmia in which ectopic firing in the atrium triggers chaotic electrical activity and rapid heartbeat. Despite their apparent dissimilarity, SAN dysfunction and AF frequently occur in the same individual^15^, and are often presumed to reflect sequalae of a shared underlying fibrotic atrial substrate.^16,17^ However, both GWAS and Mendelian Randomization have suggested a deeper mechanistic connection between SAN function and AF^18–20^, in which common variants associated with slower resting heart rate may also drive a higher incidence of AF as has been observed for a rare variant at the *PITX2* locus^21^. Determining whether slower heartbeat causes AF, or whether there are shared variants that influence both traits, could have major therapeutic consequences, but is challenging to study in genetically tractable small animal models due to species differences in heart rhythm.

Human induced pluripotent stem cells (hiPSCs) can be differentiated into various cardiomyocyte subtypes, including atrial^22^, ventricular^23^, and SAN^24–26^. PC subtypes were resolved from SAN differentiations using single cell (sc)-RNA sequencing and clustering analysis^27^, raising the possibility of using *in vitro* systems to model specific common variants associated with heart rhythm in the cell subtypes of interest, to gain insights into the pathogenesis of arrhythmia, and to identify mechanisms that mediate the connection between SND and atrial fibrillation. More fundamentally, common variants are presumed to act by affecting gene expression through cis-regulatory elements^28,29^, so a model system that permits systematic interrogation of the regulatory architecture of SAN cell types could identify transcriptional mechanism that control cell subtype specialization.

Here we used a scalable monolayer hiPSC SAN differentiation protocol to yield SAN head, SAN tail, atrial-like transition zone cells, and sinus venosus myocardium. Single cell (sc)-RNA-seq and sc-ATAC-seq defined the transcriptomic and epigenetic signatures of these cell subtypes, established their fidelity to their *in vivo* counterparts, and identified transcriptional networks engaged by different cell subtypes. Next, by using parallel reporter assays and leveraging allele-specific chromatin accessibility, we identified many novel SAN enhancers, including several that contained causative SNPs. These data provide insight into the mechanisms of trait heritability and define a role for genetically encoded autonomic responsiveness in mediating the association of sinus node function and AF.

## RESULTS

### Differentiation of Sinoatrial Node-Like Cardiomyocytes for Multi-Omics Profiling

To differentiate Sinoatrial Node-Like Cardiomyocytes (SANCMs), we modified established monolayer differentiation protocols for ventricular cardiomyocytes (VCMs) and atrial cardiomyocytes (ACMs)^30–32^ by supplementation with BMP4, inhibition of TGFβ and FGF, and lactate selection^24,33^ (Fig. 1a, Supplementary Table 1). At day 20, SANCMs beat at 2.8 +/- 0.1 Hz, ACMs at 2.1 +/- 0.1 Hz, and VCM at 1.3 +/- 0.1 Hz (n = 4 wells for each condition, Extended Data Fig. 1a). By qPCR, all CMs expressed *TNNT2*, *NKX2-5*, *ACTN2*, *MYH6, and SCN5A* (Extended Data Fig. 1b), VCMs were enriched for *MYL2*, *IRX4, MYH7* and *HEY2* ( Extended Data Fig. 1c), while ACMs were enriched for *NPPA*, *MYL7*, *NR2F2* and *KCNA5* (Extended Data Fig. 1d). Compared to VCMs, SANCMs were enriched for PC genes *TBX18*, *SHOX2*, *ISL1, HCN1*, *HCN4*, and *KCNJ3* (Extended data Fig. 1e). ^22,34^Finally, action potential waveforms for spontaneously beating cardiomyocytes showed a steeper phase 0 and prolonged phase 2 for VCMs, while SANCMs exhibited more rapid diastolic depolarization, shorter cycle length, shorter APD, and smaller AP amplitude (Figure 1b and Extended Data Fig.1f). SANCMs were also dose-dependently sensitive to Zatebradine, a specific blocker of HCN channels (Extended Data Fig 1j). At the protein level, more than 95% of cells measured in both differentiations were *TNNT2+*, (Extended Fig 2a-e), while 65%,75%, and 79% of SANCMs were SHOX2+, ISL1+, and HCN4+, respectively, as compared to 4%, 8% and 23% of VCMs (*p* < 0.0001 for each gene; Extended Fig. 2a-e). For NKX2-5, 34% of SANCMs were positive while 85% of VCMs were positive and the average signal intensity per positive nucleus was reduced by 50% in SANCMs (Extended Data Fig 2f), indicating a successful differentiation.

**Figure 1:**
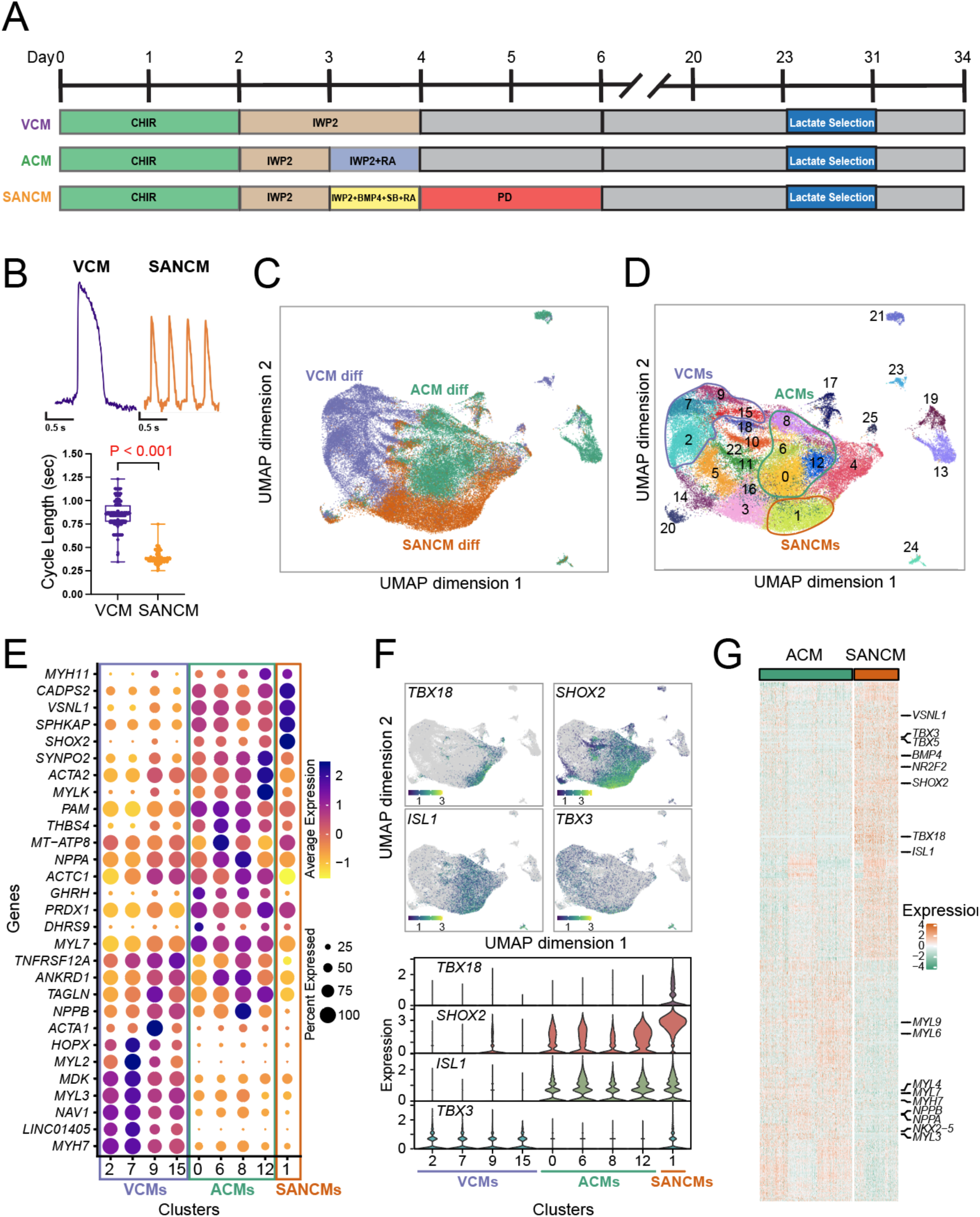
Generation and characterization of hiPSC derived Cardiomyocytes. (A) Differentiation protocols for hiPSC derived ventricular cardiomyocytes (VCM), atrial cardiomyocytes (ACM), and sinoatrial node cardiomyocytes (SANCM) (B) (*Top*) representative optical action potentials from a VCM (purple, left) and an SANCM (orange, right). (*Bottom*) panel cycle length of spontaneous beating of VCMs (n = 399) and SANCMs (n = 199) compared with a Students T-test. (C) Uniform manifold approximation and projection (UMAP) of single cell RNA sequencing data from 3 differentiations each of VCMs, ACMs, and SANCMs (n = 21,000 – 22,000 cells per condition). (D) Unsupervised clustering of the UMAP space demonstrates 26 total clusters. Based on canonical markers, circled clusters 2, 7, 9, and 15 represent VCMs, 0, 6, 8, and 12 are ACMs, and cluster 1 are SANCMs (E) Top 10 marker genes for the clusters annotated as VCM, ACM, and SANCM with expression level and percent of cells expressing each gene indicated by color and size of each dot, respectively. (F) Expression of SAN transcription factors TBX18, SHOX2, ISL1, TBX3 in UMAP space colored by expression level (*top*) and visualized by cluster in a violin plot (*bottom*). (G) Heatmap of differentially expression genes between SANCMs and ACMs recapitulate established expression differences observed between these cell types *in vivo*, including enrichment of VSNL1, TBX3, SHOX2, BMP4, and ISL1 in SAN; and enrichment of NKX2-5 and NPPA in the atrium.

### Single-Cell Transcriptomics Reveals Distinct Human Atrial, Ventricular, and SAN Populations *In Vitro*

Next, we performed single cell RNA sequencing (scRNA-seq) on day 34 VCMs, ACMs, and SANCMs from 3 independent sets of differentiations (n = 21,000 – 22,000 cells per condition, Extended Data Fig. 3a). Unsupervised clustering and dimension reduction yielded 26 clusters at a resolution of 0.6 (Fig. 1c,d and Extended Data Fig. 3b-d). Cells derived from VCM, ACM, and SANCM directed differentiations occupied distinct regions in the UMAP space with minimal overlap and were similar for each replicate, indicating unique expression signatures (Fig. 1c, Extended Data Fig. 3e). The largest number of cells represented *TNNT2^+^, ACTN2^+^,* and *MYH6^+^*cardiomyocytes (Fig. 1d). Using a cell composition cut-off of 70%, 8, 3 and 6 clusters originated from the ACM, SANCM, and VCM differentiation, respectively, with the remaining clusters representing cells derived from multiple differentiations (Extended Data Fig. 4a).

Individual clusters were assigned to cardiomyocyte subtypes by intersecting their marker genes with established expression patterns (Fig. 1d,e; Extended Data Fig. 4b; Supplementary Table 2). Clusters 2, 7, 9, 15 represent VCMs (*MYL2, MYH7, HEY2, IRX4,* and *HOPX*), clusters 0, 6, 8 and 12 represent ACMs (*MYL7, NPPA, NR2F2. KCNJ3,* and *KCNA5*) and cluster 1 represents PCs (*SHOX2, ISL1, TBX18,* and *TBX3*). We also identified several non-cardiomyocyte clusters (C10, C13, C14, C17, C19, C20, C21, C23, C24 and C25). that included proepicardial cells (C25) marked by *WT1, ALDH1A2, BNC1,* and *PDPN*, endocardial cells (C13, C19) marked by *FOXC1, MSX1, NRG1, NPR3* and *NFATC1*, primitive mesendoderm marked by *FOXA2, EOMES, MESP1* and *MESP2* (C17, C21), and visceral neurons marked by *PHOX2B, PRPH,* and *CHRNA3* (C24) (Fig. 2b,c and Extended Data Fig. 4c,d)

**Figure 2:**
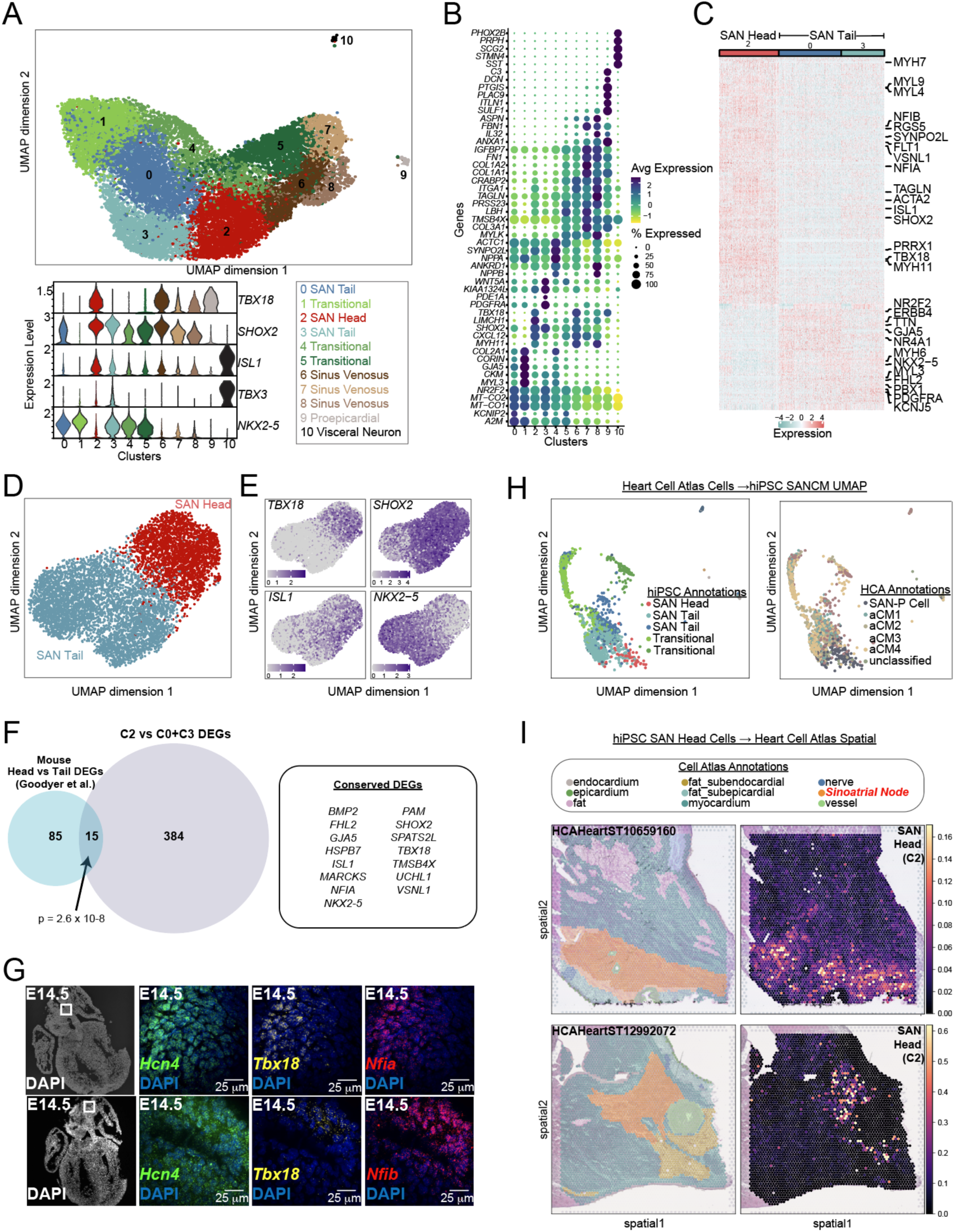
HiPSC derived SANCMs Recapitulate Cellular Diversity of the SAN. (A) Uniform manifold approximation and projection (UMAP) of scRNA-seq on day 34 hiPSC-derived SANCMs after unsupervised clustering defined 10 clusters, annotated below with SAN subtype labels (below, right) that are based on expression of key SAN and atrial transcription factors *TBX18, SHOX2, ISL1, TBX3,* and *NKX2-5*, as shown in the violin plot (below, left). (B) Dot plot of top 5 marker genes that were observed in each cluster with expression level and percent of cells expressing each gene indicated by color and size of each dot, respectively. (C) Heatmap of differentially expressed genes between SAN Head cells (cluster 2) and SAN Tail cells (clusters 0 and 3 combined) demonstrates enrichment of *ISL1, TBX18, VSNL1, NFIA* in SAN Head and enrichment of *NKX2-5, NYH6* and *TTN* in San Tail cells. (D) UMAP of sub-clustered SAN Head cells (cluster 2) and SAN Tail cells (clusters 0 and 3) only shows clear spatial segregation of the two cell types with minimal intermixing, while (E) feature plots demonstrate complementary expression of *TBX18* and *NKX2-5* in SAN head and tail, with a gradient of *ISL1* and *SHOX2* expression from SAN head to SAN tail. (F) Venn diagram showing the overlap of conserved differentially expressed genes (DEGs) from Human iPSC derived SAN Head vs Tail cells (our data) and Mouse *in vivo* isolated SAN Head vs Tail cells (Goodyear et al.) Fisher’s exact test was used to test for significance of overlap. (G) 3-color hybridization chain reaction fluorescent in-situ hybridization (HCR-FISH) for Hcn4, Tbx18, and Nfia (*top*) and Hcn4, Tbx18 and Nfib (*bottom*) in embryonic day 14.5 mouse heart sections, focusing on the SAN Head. The left panels show staining of the entire section with 4’,6-diamidino-2-phenylindole (DAPI) and the white square indicates the area of detail shown on the images on the right. *Nfia* and *Nfib* colocalize with *Tbx18* in the San head region while *Hcn4* is expressed throughout the SAN. (H) Projection of human SAN cells from the Heart Cell Atlas (HCA) into the UMAP shown in panel A with SANCM annotations (left panel) and HCA annotations (right panel), demonstrating that human adult pacemaker cells (SAN-P cell) map most closely to cluster 2 (SAN head) from our dataset, while different atrial cell subtypes map to SAN tail and transitional cells. (I) Annotated tissue sections from the SAN regions of 2 adult human subjects (1 subject per row) from the HCA with cell type annotations indicated by color on the left (SAN region in orange). Right panels show a projection of hiPSC-SANCM cluster 2 into the tissue plane with most SAN head cells mapping into the histological central portion of the adult SAN.

Differential expression analysis between SANCMs and ACMs yielded 506 genes (DEGs) that recapitulated established differences between atrial and PCs such as upstream transcriptional regulators (TBX18, TBX3, NKX2-5, SHOX2, and ISL1)^35^ (Fig. 1g, Supplementary Table 3). These DEGs were associated with biological processes related to sinoatrial node development, cardiac conduction, and regulation of heart contraction (Supplementary Table 4). Intersecting the list of DEGs with neonatal mouse^36^ and human fetal^37^ SAN versus right atrium comparative transcriptomic datasets demonstrated significant overlap among the three datasets (Extended Data Fig. 4e *p* = 0.02 for overlap between SANCMs and human dataset and *p* = 6.4 × 10^−7^ for overlap between SANCMs and mouse dataset).

### Sub-clustering of SAN Differentiation Identifies Specialized PC Subtypes

The functional organization of the SAN, diagrammed in Extended Data Fig. 5a, includes a more autonomically responsive SAN head located cranially, an SAN tail extending inferiorly along the venous valves, and a transition zone with more atrial-like SAN myocytes that facilitates source-sink matching between SAN and atrium^7^. Different PC subtypes express different levels of key transcriptional regulators that guide their distinctive phenotypes^8^ (Extended Data Fig. 5b).

To test whether discrete PC subtypes could be identified within our SANCM population, we further sub-clustered on SANCMs, yielding 10 subclusters (Fig. 2a,b; Extended Data Fig. 5c-f). Cluster 2 resembled SAN head cells (*TBX18*^HI^/SHOX2^+^/ISL1^HI^*/TBX3*^+^/*NKX2-5*^LOW^), clusters 0 and 3 resembled SAN tail cells (TBX18^−^/SHOX2^+^/ISL1^LOW^/TBX3^+^/NKX2-5^+^), clusters 1 and 4 were transitional cells (*TBX18*^−^/SHOX2^+^/ISL1^−^*/TBX3*^LOW^/*NKX2-5*^HI^), and clusters 6, 7, and 8 represented sinus venosus myocardium (*TBX18*^+^/SHOX2^+^/ISL1^LOW^*/TBX3*^−^/*NKX2-5*^LOW^). The two SAN tail cell clusters were distinguished by differential expression of the SAN transcriptional repressor *TBX3* and its downstream target genes such as *NPPA*, possibly reflecting residual heterogeneity within these populations or varying degrees of maturation (Fig. 2a, Extended Data Fig. 5g). Notably, cluster 5 most resembled transitional myocardium (*NKX2-5*^HI^) but expressed some markers of sinus venosus myocardium including *TAGLN* at high level. We identified small populations of proepicardial (cluster 9) and visceral neurons (cluster 10); excluding these non-myocyte clusters did not appreciably affect the overall clustering of cell types (Extended Data Fig.5h).

Direct comparison identified several hundred DEGs between SAN head (C2) and SAN tail (C0+C3) populations, encompassing GO terms for biological processes related to cellular morphogenesis, cytoskeletal organization, and cell adhesion. Supporting the autonomic response function of the SAN head, GO terms enriched in SAN head included neuron protrusion/axon development and synapse formation (Fig. 2c and Supplementary Table 5). SAN Head and Tail cells had contrasting expression of *TBX18* and *NKX2-5,* with greater expression of *SHOX2* and *ISL1* in SAN head versus SAN tail cells (Fig. 2d,e).

Importantly, 15 out of 85 mouse SAN head vs tail DEGs identified in a recent study^9^ were also differentially expressed between C2 (SAN head-like) and C0+C3 (SAN tail-like), an odds ratio of 7.8 over SAN-expressed genes that were not on the mouse head versus tail DEG list (*p* = 2.6 × 10^−8^ by Fisher’s exact test, Fig. 2f). The list of shared DEGs between mouse and human included canonical transcription factors *ISL1, TBX18, SHOX2* and *NKX2-5*, supporting the existence of an evolutionarily conserved blueprint not only for SAN development broadly but also for fine patterning of PC subtypes. Other SAN Head vs Tail DEGs conserved between mouse and human included *GJA1*, an established downstream target of *NKX2-5* (enriched in SAN tail), modulators of signal transduction (*FHL2*, *VSNL1*, *BMP2*), and cytoskeletal organizing proteins (*MARCKS, TMSB4X*). We also observed conserved enrichment in SAN Head cells for Nuclear Factor I A (*NFIA)*, a transcription factor without a previously known role in SAN development. Expression of *Nfia* and *Nfib* in murine E14.5 SAN measured by in situ hybridization showed colocalization with *Hcn4*+ and *Tbx18*+ in the SAN head region, providing addition support for their evolutionarily conserved expression patterns (Fig. 2g).

To test for concordance with other SAN differentiation methodologies, we projected scRNA-seq data from a published hiPSC SAN differentiation^27^ into our UMAP and found strong agreement in cluster assignments between the two protocols, such that 94% of SAN head cells mapped to either our SAN head cluster (C2) or to the narrow interface between our SAN head and the nearest SAN tail clusters (C2/C3 interface); 90% of SAN tail cells mapped to our SAN tail clusters (C0+C3); 82% of transitional cells mapped to our transitional clusters (C1+C4), 64% of sinus venosus cells mapped to our sinus venosus clusters (C5+C6+C7+C8), and 97% of proepicardial cells mapped our proepicardial cluster (C9) (Extended Data Fig 5i). To test for concordance with primary PCs, we projected cells from a single cell multi-omics study of adult human SAN tissue^38^ into the SANCM UMAP space. In this analysis, 78% of human primary PCs (192/245) were assigned to SAN head or SAN tail, while 18% were assigned to transitional cell clusters. In contrast, only 9% of primary atrial cardiomyocytes were classified as SAN head or SAN tail, of which just 0.002% (18/9595) were assigned SAN head identity. We also performed the converse mapping, in which we projected our SAN head cells (C2) into annotated human tissue sections from 3 donor hearts. In each heart, our SAN head cells mapped most closely to the central SAN region surrounding the nodal artery (Fig 2i). Taken together, the concordance of our SAN head versus tail differential expression analysis with findings from primary SAN tissues, and with other hiPSC-based SAN datasets, strongly supports the fidelity of our system to model specialized cell types in the cardiac inflow and venous pole.

### scATAC-seq Identifies Cell Type Specific Regulatory Regions in SANCMs

To define cell subtype-specific epigenetic signatures, we performed single cell assay for transposase accessible chromatin with sequencing (scATAC-seq) on cells from 3 independent SANCM differentiations. UMAP projection identified 8 clusters (Fig. 3a and Extended Data Fig. 6a) which we annotated based on predicted expression of canonical SANCM marker genes: *ISL1^+^/TBX18^+^/SHOX2^+^/NKX2.5^LOW^* SAN head (C2), *ISL1^+^/SHOX2^+^/NKX2.5^+^/TBX18^LOW^* SAN tail (C1, C4, C6), *NKX2.5^HIGH^/SHOX2^LOW^/TBX18^−^* transitional cells (C0, C5), and TBX18+/NKX2-5- sinus venosus cells (C3) (Fig. 3b, Extended Data Fig. 6b,c). Re-clustering of the SAN head and SAN tail cells alone demonstrated expected patterns of predicted gene expression for canonical SAN transcriptional regulators, accurately mirroring transcriptional start site permissiveness (Extended Data Fig. 6d).

**Figure 3.**
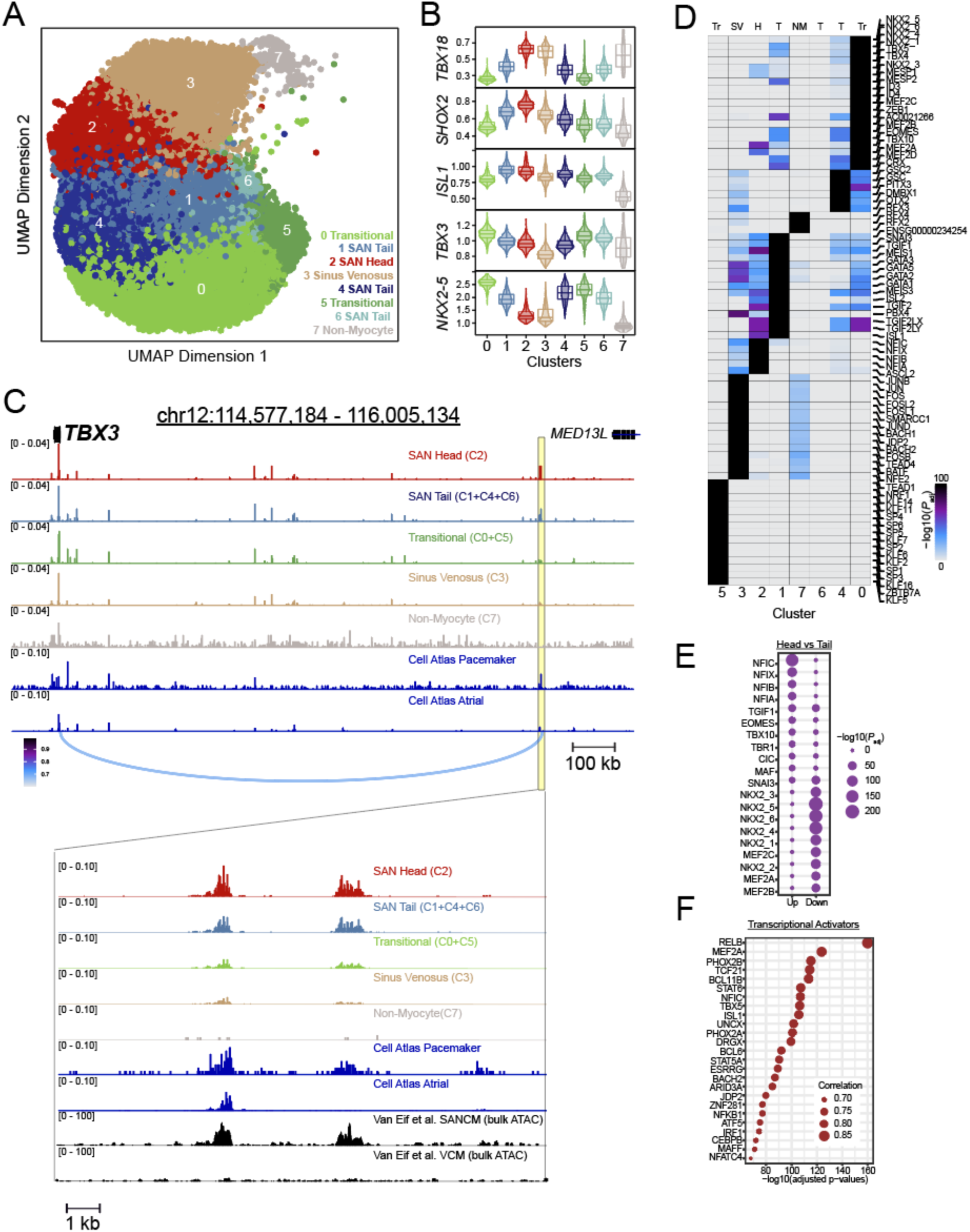
Single Cell ATAC-seq of Day 34 hiPSC derived SANCMs. (A) Uniform manifold approximation and projection (UMAP) performed on single cell ATAC-seq data from 3 replicates of day 34 SANCMs with cluster annotations for pacemaker cell subtypes, preserving the cell subtype color scheme used for annotation of scRNA-seq data shown in Figure 2. (B) Predicted expression patterns of canonical SANCM marker genes based on promoter region chromatin accessibility for each SAN transcription factor. (C) *Top panel:* Chromatin accessibility tracks at the TBX3 locus from SANCM scATAC-seq re-bulked by cluster annotations, with re-bulked scATAC-seq tracks from Heart Cell Atlas pacemaker cells and atrial cardiomyocytes below. Peak-to-gene (P2G) links defined by applying ArchR to scATAC-seq and scRNA-seq are shown at the bottom with the color scale representing correlation strength. A single P2G link was observed for an accessible chromatin area (highlighted in yellow) approximately 1-Mb from TBX3 at an established TBX3 SAN enhancer. *Bottom panel:* Chromatin accessibility tracks for the region highlighted in yellow in the top panel, with addition of bulk ATAC-seq data from SANCMs and VCMs from a prior study are shown below (Van Eif et al). C2 (SAN head cells) exhibit the highest accessibility at the *TBX3* SAN enhancer followed by SAN tail cells (C1+C4+C6), with lower accessibility for other clusters. Peak heights were normalized and scaled by the ReadsInTSS parameter, the default normalization method in ArchR (D) Hypergeometric enrichment of top 20 transcription factor binding motifs in marker peaks of each cluster, colored by corrected P value. SAN head cells marker peaks(C2) were enriched for *ISL1* and *NFI* sites, while marker peaks for the more atrial-like C0 (transitional cell) showed enrichment of *NKX2-5*, *MEF2*, and *TBX* factors. (E) Top 10 upregulated and downregulated transcription factor binding motifs in SAN Head cells (C2) versus SAN Tail cells (C1+C4+C6). (F) The 25 transcription factors whose expression is most strongly correlated to increased accessibility of their corresponding motif and shown in order of statistical significance.

Next, we integrated our scATAC-seq and scRNA-seq datasets of SANCMs by assigning RNA-seq profiles to each scATAC-seq cell using canonical correlation analysis (CCA) and we subsequently identified candidate regulatory elements highly correlated to RNA expression of proximal genes (Supplementary Table 6), many of which supported distal enhancer-promoter links of up to several hundred kilobases, along with quantitative estimation of interaction strengths. Genomic tracks for accessibility, marker peaks and peak-to-gene links in different SANCM subtypes are shown for *TBX3*, which has a well-established SAN enhancer lying nearly a megabase away from the transcriptional start site that was the only peak-to-gene link observed (Fig. 3c)^39^. A pseudobulk of scATAC-seq data corresponding to PCs and ACMs from the Human Cell Atlas project demonstrated persistence and differential accessibility of this region in adult human hearts, suggesting that cis-regulation of *TBX3* expression may be established in the embryo and maintained throughout adulthood. Comparison with another dataset ^39^ on a mixed SANCM population revealed that most of the ATAC-seq signal were derived from SAN head cells, with some contribution from SAN tail cells, suggesting that the upstream processes that regulate accessibility of the *TBX3* enhancer may regulate fine-patterning of the SAN (Fig. 3c, bottom panel).

### Identification of Transcriptional Pathways Important for SAN Subtype Differentiation

To identify upstream regulators that drive SAN subtype-specific differentiation, we performed motif enrichment analysis of marker peaks for each cluster (Fig. 3d, Extended Data Figure 3f, and Supplementary Table 7). We found that MEF2, TBX5, and NKX2-5 sites were enriched in transitional cell peaks, whereas marker peaks for the largest SAN Tail cluster (C1) demonstrated enrichment of ISL and GATA binding sites, as well as sites for the TALE-class transcription factors MEIS and TGIF. The SAN Head cluster (C2) exhibited the strongest enrichment for the NFI sites, followed by ISL1, while the sinus venosus cluster (C3) demonstrated strong enrichment for AP-1 sites. These findings were underscored by a direct comparison of motif enrichment between SAN Head and SAN Tail marker peaks, in which NFI sites were strongly enriched in SAN head whereas motifs for cardiomyogenic factors NKX2-5 and MEF2 were enriched in SAN tail (Fig. 3e and Supplementary Table 8). Motif enrichment analysis for the subset of ATAC-seq peaks that were positively linked to transcription via peak-to-gene analysis identified the top 20 positive transcriptional regulators across the dataset (Fig. 3f). This analysis identified known regulators of SAN development ISL1, TBX5, and MEF2 but also identified NFI motifs, which exhibit cellular subtype-specific motif enrichment and differential gene expression (NFIA) in SAN head versus other venous pole cardiomyocytes.

### Chromatin Accessibility Regions Demonstrate Enhancer Activity in SANCMs

Regions of differential accessibility between SAN subpopulations were also observed near *TBX18, ISL1, SHOX2, and NKX2-5*, including at established and novel cis-regulatory elements, many of which exhibited differential accessibility in pseudobulked scATAC-seq data from PCs and ACMs from the Heart Cell Atlas.^36,39^ (Fig. 4a-d, Extended Data Fig. 7a,b) In addition, numerous cluster-enriched ATAC-seq peaks were identified (Extended Data Fig. 6c). For example, the syntenic region to a mouse Isl1 enhancer previously characterized by our group demonstrated differential accessibility in PCs (SAN head and tail) as compared to more atrial-like transitional cells (Fig. 4a, element designated ‘ISL1-R3’), analogous to ISL1 mRNA levels in these subpopulations. In addition, our analysis identified other ISL1 enhancers linked to transcription that were previously validated in a zebrafish system (ISL1-R1/R2) ^39^. When ISL1-R3 and ISL1-R2 elements were experimentally validated via a luciferase reporter system, both elements robustly increased luciferase activity in SANCMs while *suppressing* basal promoter activity in VCMs (Fig. 4b, Extended Data Fig. 7e). Similar results were obtained with established enhancers for SHOX2^39^ (Extended Data Fig. 7a,c) In contrast, when we tested 2 open chromatin regions at the *NXK2-5* locus, both demonstrated significantly *reduced* luciferase activity as compared to the empty vector, suggesting that these elements may be repressive (Extended Data Fig, 7b,d).

**Figure 4.**
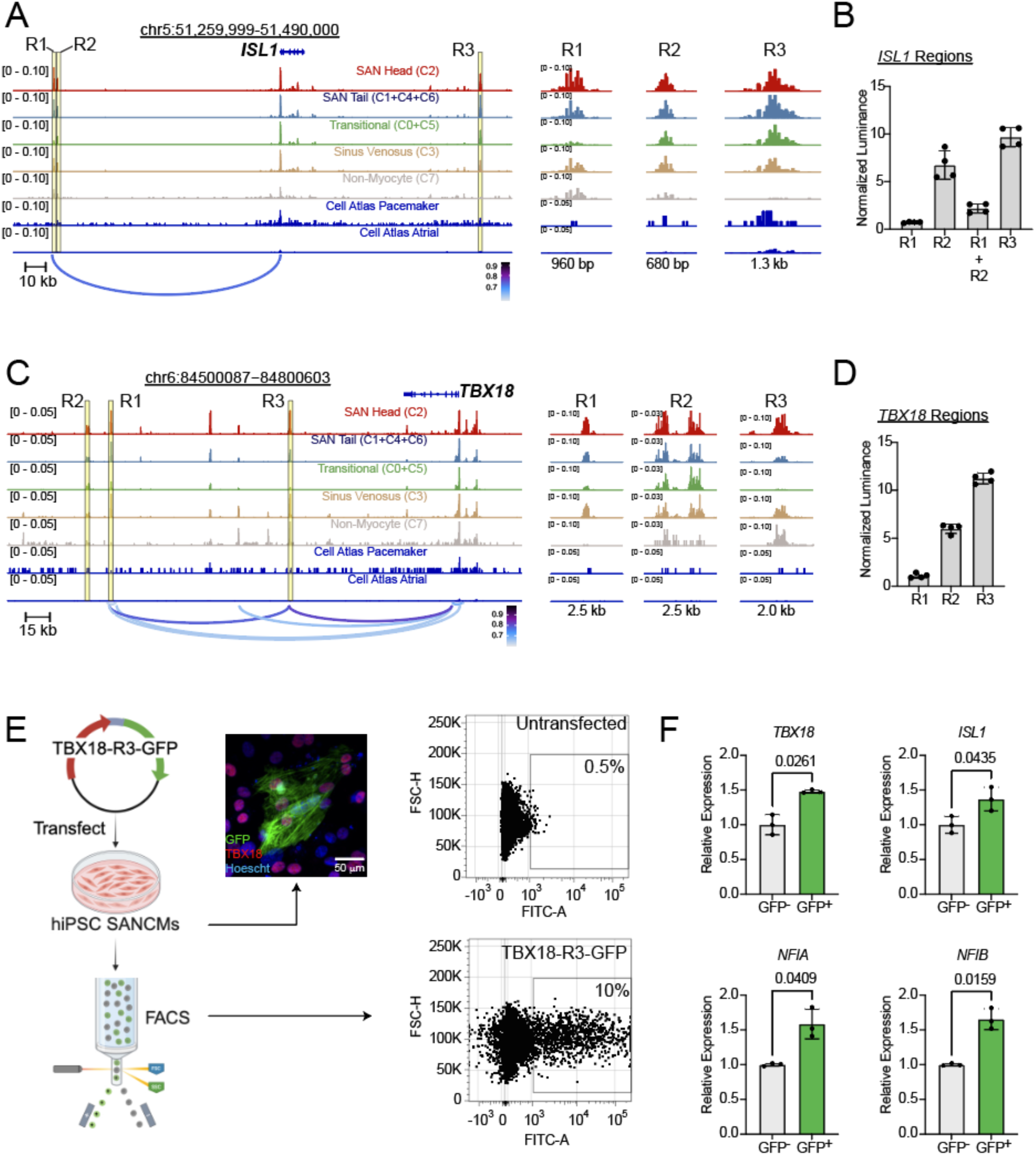
Characterization of SANCM Accessible Chromatin Regions at ISL1 and TBX18 Loci. (A) Chromatin accessibility tracks at the *ISL1* locus from SANCM scATAC-seq re-bulked by cluster annotations, with re-bulked scATAC-seq tracks from Heart Cell Atlas pacemaker cells and atrial cardiomyocytes below. Peak-to-gene (P2G) links defined by applying ArchR to SANCM scATAC-seq and scRNA-seq are shown at the bottom with the color scale representing correlation strength. Peak heights were normalized and scaled by the ReadsInTSS parameter, the default normalization method in ArchR. Three accessible chromatin regions (R1, R2, and R3) are highlighted in yellow and shown at higher magnification in the panels to the right. R3 is a previously identified ISL1 enhancer (Galang et al, 2020). (B) Luciferase/Renilla reporter activity for 4 plasmid constructs generated for R1, R2, R1+R2, and R3 are shown normalized by activity of the empty luciferase vector in transfected SANCMs (n = 4 biological replicates). *P* values are shown for comparison to a value of 1 corresponding to empty vector. (C) Browser tracks analogous to panel A that highlight three regulatory elements at the TBX18 locus that were cloned for functional analysis. Element TBX18-R3 and TBX18-R1 have differential accessibility in SAN head over SAN tail and transitional cells. (D) Luciferase/Renilla activity for the three TBX18 locus putative regulatory elements displayed as fold-change over empty vector (*n* = 4 transfections for each plasmid). P values denote comparison to a value of 1 as in (B). (E) (Left) Experimental flowsheet (Biorender) for transfection of TBX18-R3-GFP into hiPSC-SANCMs followed by immunostaining to visualize overlap of GFP and TBX18 expression (middle), and isolation of GFP+ cells and GFP- cells using FACS (Right). Comparison of untransfected cells (top, right) with transfected cells (bottom, rights) demonstrates that GFP+ comprise about 10% of the total SANCM culture. (F) Quantitative PCR for SAN head enriched genes *TBX18*, *ISL1*, *NFIA*, and *NFIB* demonstrates enrichment for all 4 transcripts in TBX18-R3-GFP+ cells (n = 3 transfections, statistical comparison with t-test with raw P values indicated).

Despite its importance for SAN formation and gene regulation, specific regulatory elements controlling TBX18 expression in the SAN have not yet been defined. In our data, the TBX18 locus demonstrated differentially accessible regions in different SAN subtypes (Fig.4c). When three of these elements were experimentally validated, the element demonstrating the most robust cell-type differential accessibility in SAN head and sinus venosus cells (TBX-R3) was also the element that demonstrated the most robust luciferase activation (Fig. 4d). To test whether this element could exhibit differential activity across different SAN cell subtypes, we generated a TBX18-R3-GFP enhancer-reporter plasmid which we transfected into SANCMs for fluorescence activated cell sorting (Fig 4e), demonstrating that about 10% of the SANCMs were robustly GFP+. Furthermore, purified TBX18-R3-GFP cells exhibited significantly higher expression of several transcripts that are differentially expressed in SAN head cells, including *TBX18*, *ISL1*, *NFIA*, and *NFIB* (Fig. 4f). Taken together, these data provide strong experimental evidence that novel regulatory elements discovered in our scATAC-seq dataset are functional enhancers with differential activity across SAN cell subtypes.

### Integration of Multi-Omics Data with Human Genetic Data for Gene Discovery and Annotation

Because cell types identified in SANCM cultures have transcriptional and epigenetic signatures that resemble their adult human counterparts, we hypothesized that putative regulatory regions in SANCM subtypes are enriched for single nucleotide polymorphisms (SNPs) associated with heart rhythm-related traits. To test this hypothesis, we recreated Manhattan plots using summary statistics from previously published GWAS for sinus node function^40,41^ and for atrial fibrillation^42,43^ but restricted the statistical analysis only to regions of open chromatin in SANCM clusters (C0, C1, C2, C4, and C6 in Fig.3a) identified from our scATAC-seq dataset (Fig. 5a). This restricted analysis recapitulated most of the genetic associations identified in the source studies even when restricted to marker peaks for SAN head, SAN tail, and transitional cells (Extended Data Fig. 8a,b; Supplementary Table 9), supporting the relevance of our putative regulatory elements to genetic variation of heart rhythm in human populations.

**Figure 5:**
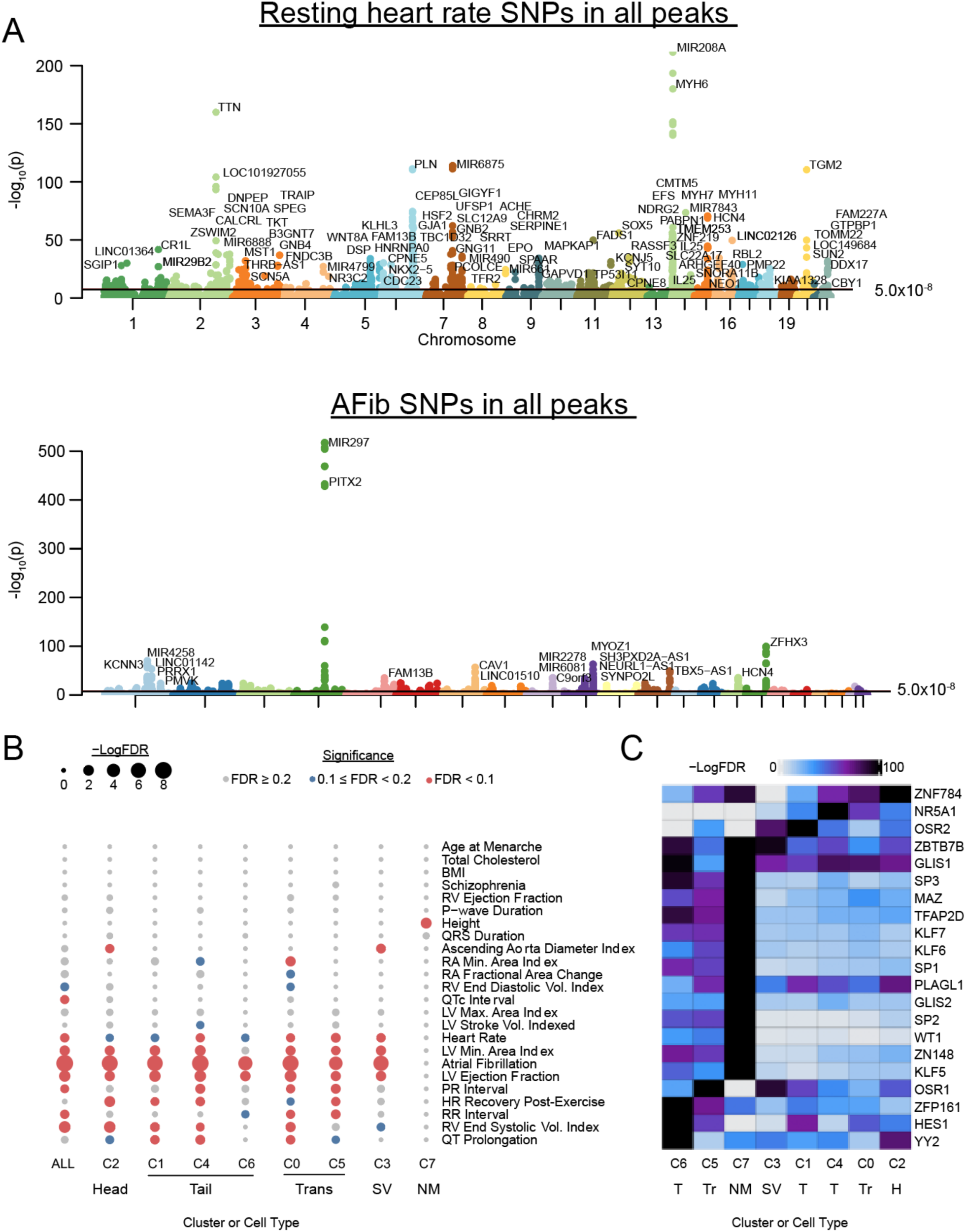
Integrative analysis of scATAC-seq data and GWAS. (A) Manhattan plots showing association of resting heart rate (top) and atrial fibrillation (bottom) SNPs ascertained from the UK Biobank cohort within all ATAC-seq peaks from our study. (B) Dot plot showing association of SAN clusters and cell types with complex traits and diseases. Cell-type-stratified linkage disequilibrium score regression (LDSC) analysis was performed using GWAS summary statistics for 41 phenotypes. All ATAC peaks identified from each fetal and adult cell type by human enhancer atlas (http://catlas.org/humanenhancer) were used as background for analysis. P-values were corrected using the Benjamini-Hochberg procedure for multiple tests. FDRs of LDSC coefficient are displayed. (C) Motif enrichment analysis by cluster for scATAC peaks containing SNPs associated with atrial fibrillation (P < 5 × 10^−8^) demonstrated distinct motif enrichment patterns in different cell types. Abbreviations: Head (H), SAN Head; Tail (T), SAN Tail; Trans (Tr), Transitional; SV, Sinus venosus; NM, non-myocyte; ALL, pseudo-bulk of all clusters.

Next, we applied stratified linkage disequilibrium score regression (LDSC) to evaluate whether heritability for heart rhythm-related traits, general cardiovascular traits, or selected non-cardiac traits was enriched in candidate regulatory elements from SANCM cell subtypes (Fig. 5b, Extended Data Fig. 8c). As expected, traits that are not directly related to cardiac physiology failed to show significant associations with regulatory regions of SANCM cells whereas multiple cardiac structural and functional traits showed differing degrees of association. Heritability of heart rate and RR interval was more significantly enriched within SNPs in the peaks of SAN tail and transitional cells than SAN head, while heart rate recovery after exercise (an indicator of cardiac autonomic responsiveness) was more significantly associated with SAN head regions, highlighting the concordance of our cluster assignments with *in vivo* functional classifications of these cells. Furthermore, accessible chromatin regions were enriched for determinants of atrial fibrillation risk to differing degrees in all cardiomyocyte cell types but more strongly than for any other cardiac or non-cardiac traits including left ventricular ejection fraction, right ventricular function, PR interval and QT interval, revealing that regulatory regions identified in SANCMs contain common variants that influence both SAN function and AF risk.

We also performed motif enrichment analysis by cluster for scATAC-seq peaks containing SNPs with genome-wide significance for atrial fibrillation (Fig. 5C). This analysis demonstrated that different scATAC-seq clusters exhibited distinct patterns of motif enrichment, with non-myocyte cells (C7, not significant in LDSC analysis) having a set of unique TF motifs in AF-associated sites including *WT1*, a regulator of epicardial formation, injury response, and a contributor to atrial fibrillation pathogenesis. ^44,45^ Within the SAN clusters (most prominently in SAN tail cluster C1 but also in sinus venosus (C3) and SAN head (C2)), we observed enrichment of motifs for odd-skipped-related-1 (*OSR-1*), a posterior second heart field transcription factor required for atrial and venous pole development.^46–48^

### Functional Assessment of Putative Trait-Associated Enhancers

The association of open chromatin regions with heritability of heart rate and atrial fibrillation motivated us to pursue functional studies to identify trait-associated enhancers and to define the effects of common variants on enhancer activity. To test whether scATAC-seq peaks containing trait-associated SNPs could function as enhancers, we used self-transcribing active regulatory region sequencing (STARR-seq)^49^, a parallel reporter assay in which active enhancers self-transcribe, allowing RNA sequencing to quantitatively report enhancer activity (Fig. 6a). For this assay, we selected 331 500-bp candidate regulatory elements that were repeatedly observed with a peak score of above 200, each containing at least one SNP associated with either sinus node function (n = 224) or AF (n = 107). 5 genomic regions were selected with no nearby ATAC-seq peak to serve as negative controls for the assay (Fig. 6a, Supplementary Table 10). For each peak, we included a reference and alternative allele to test whether allelic imbalance could be observed for any of the trait-associated SNPs, an approach that has been previously termed ‘snpSTARR-seq’.^50^

**Figure 6.**
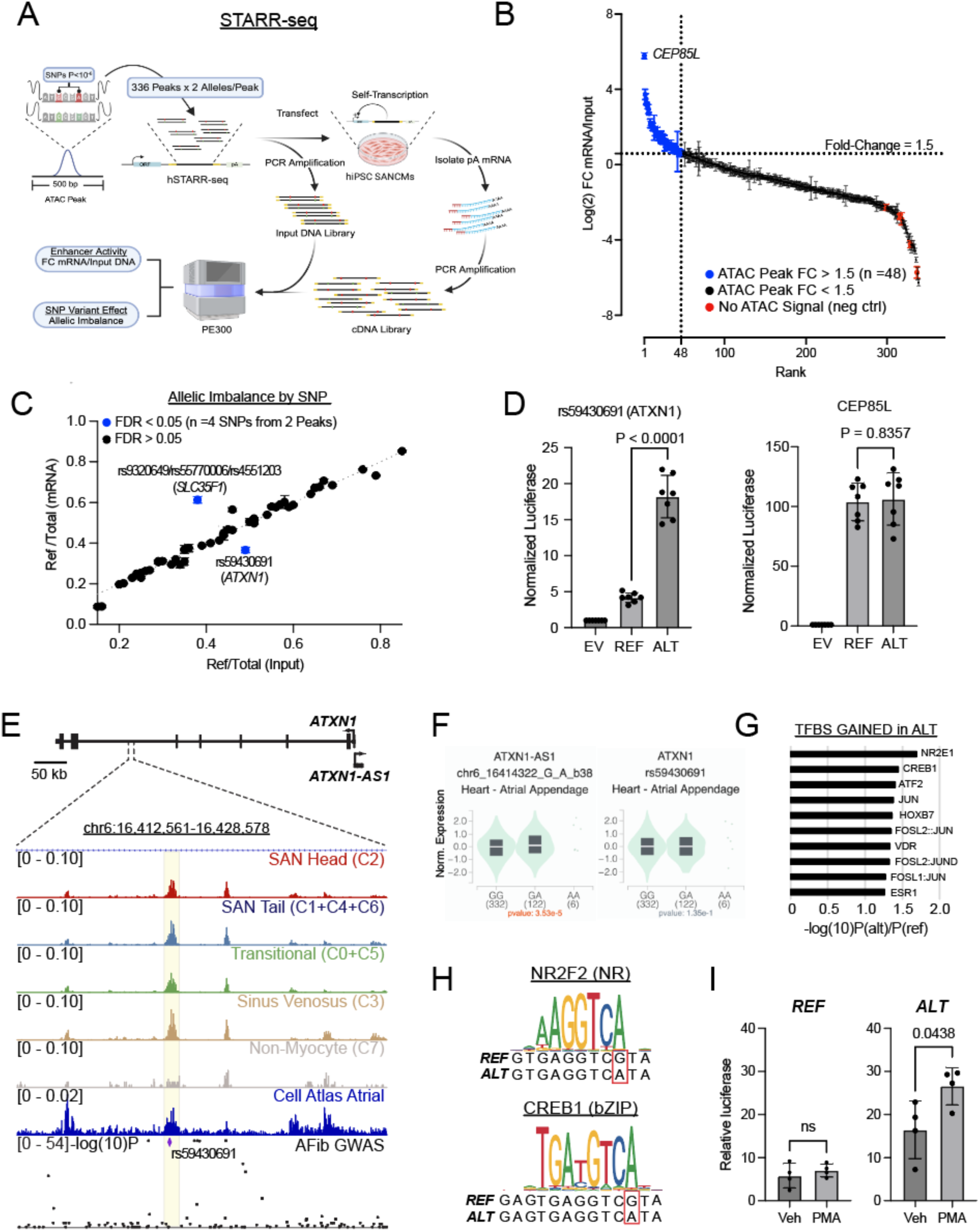
Parallel Reporter Assay in SANCMs Using STARR-Seq. (A) Experimental diagram (Biorender) for self-transcribing active regulatory region sequencing (STARR-seq) using SANCMs. 331 ATAC-seq peaks (and 5 control regions) were selected that contained a SNP associated with either resting heart rate, atrial fibrillation, or heart rate recovery after exercise. (B) Results of STARR-seq plotted as fold change enrichment of mRNA for each of 336 reference alleles ranked from highest to lowest, with 48 peaks exceeding arbitrary cutoff of 1.5 (C) Allelic imbalance between reference and alternate alleles assessed for single nucleotide polymorphisms (SNPs) occurring in the top 48 scATAC-seq peaks highlighted in (B), displayed as a comparison between the ratio of reference allele/total allele in the input cDNA to the self-transcribed mRNA. Blue denotes statistically significant allelic imbalance. (D) Luciferase/Renilla comparing the reference and alternative alleles for the SNP at (left panel) the putative ATXN1 locus and at the CEP85L locus (right) (n = 3 separate transfections on 7 distinct differentiations). P values were derived from unpaired T-tests. (E) Browser tracks highlighting the scATAC-seq peak at the *ATXN1* locus (yellow) in relation genome wide association study for atrial fibrillation, with rs59430691 highlighted as the lead SNP. Peak heights were normalized and scaled by the ReadsInTSS parameter, the default normalization method in ArchR (F) Expression of *ATXN1-AS1* and *ATXN1* in human heart samples by rs59430691(G/A) genotype from the GTEx database. (G) List of transcription factor binding sites gained with substitution of the alternate allele, defined as those with a greater than 10-fold reduction in the P value for a sequence-to-motif match. (H) Transcription factor binding motif logos for nuclear receptor family factor NR2F2 (JASPAR MA1111.1, above) and bZIP family factor CREB1 (JASPAR MA0018.4, below) with the REF and ALT sequences at rs59430691 highlighted (red box), (I) Luciferase/Renilla, normalized to an empty vector, comparing control (veh) to transient exposure to 5 nM phorbol 12-myristate 13-acetate (PMA) for the two constructs.

For each sequence tested, we quantified the fold change of mRNA over input DNA using 300-bp paired-end reads permitting accurate calling of reference and alternative alleles (Fig. 6b). As expected, we found that the 5 negative control regions were among the group with the lowest enrichment scores. Using an arbitrary enrichment score cutoff of 1.5-fold, we identified 48 regulatory elements with robust transcriptional activity in SANCMs, although most candidates we tested self-transcribed significantly more than the negative controls, consistent with some degree of enhancer activity (Supplementary Table 11).

Most genes adjacent to candidate regulatory elements were expressed in all SANCM cell types, although a few were enriched in specific clusters (Extended Data Fig. 9). While many of these enhancers were located near genes with established roles in heart rhythm regulation or in heart function, including ACHE, CAV2, AKAP6, TRPM4, and MEF2D, some occurred at loci containing genes annotated as GWAS hits for heart rate or AF but without well-established roles in cardiac physiology (e.g., ATXN1).

### A SNP at the ATNX1 Locus Confers Signal-Responsiveness on an Enhancer

For each self-transcribing enhancer included in the assay, we ascertained the fraction of mRNA reads arising from reference allele and compared this to the fraction of reference alleles for that enhancer in the input DNA library. Among our top 48 self-transcribing enhancers, 2 enhancers exhibited were deemed to exhibit allelic imbalance when these values diverged significantly (Fig. 6c), one with a single AF-associated SNP and one with 3 HR-associated SNPs. One of these enhancers is located at the *ATXN1* locus, a gene expressed in all SAN cardiomyocyte subtypes, and contains rs59430691(G/A), a SNP for which the reference allele is associated with a higher risk of atrial fibrillation and, in our assay, weaker enhancer activity. To further quantify the difference in enhancer activity, we experimentally validated the activities of the reference vs alternate alleles in a reporter assay and showed a 4-fold increase in enhancer activity with the alternate allele (Fig. 6d). As a negative control, we also tested the reference and alternate allele for the strongest enhancer identified in our STARR-seq assay at the CEP85L locus which did not exhibit allelic imbalance. As expected, this ATAC-seq peak functioned as strong enhancer in our luciferase assay with no significant difference in activity between the reference and alternate allele.

Rs59430691(G/A) is the lead SNP at the *ATXN1* genomic locus, and the associated ATAC-seq peak is accessible in all cardiomyocyte cell types including adult atrial cardiomyocytes from the Human Cell Atlas (Fig. 6e). In human atrial tissue samples from the GTEx^51,52^ database, the alternate (A) allele is associated with a trend towards increased expression of *ATXN1* and is highly significantly associated with expression of *ATXN1-AS1* whose proximal promoter region is coextensive with that of *ATXN1* (Fig. 6f). When we searched for transcription factor binding motifs changed by the SNP, we found that the alternate allele created a consensus nuclear receptor half-site (AGGTCA) and increased similarity to multiple CREB-related family motifs of the basic leucine zipper (bZIP) class, including ATF2, CREB1, and JUN-family transcription factors (Fig. 6g,h). While the nuclear receptor site could account for increased constitutive enhancer activity (for example, via the atrial factor *NR2F2*), we wondered whether the alternate allele might confer signal responsiveness as well via the bZIP neo-site or a signal responsive nuclear receptor. To test this possibility, we repeated the luciferase assay in the presence of vehicle or 5 nM phorbol 12-myristate 13-acetate (PMA), which activates protein kinase C (PKC)-dependent signaling pathways and induces AP-1, ATF, CREB, and NR4A transcriptional programs (Fig. 6i). Whereas the *ATXN1* enhancer with the reference allele did not exhibit a significant change in luciferase activity after 5 nM PMA exposure, the *ATXN1* enhancer bearing the alternate allele responded to the PMA with a significant increase in luciferase activity, suggesting that the alternate allele of rs59430691 may be protective against AF through constitutive and stress-induced induction of *ATXN1* expression.

### Chromatin Accessibility Allelic Imbalance

To expand our analysis beyond the 331 variants we evaluated by STARR-seq, we investigated whether we could prioritize likely causal variants by searching for chromatin accessibility allelic imbalance (CAAI) at genomic loci that were heterozygous for trait-associated SNPs occurring within our candidate regulatory elements. We first performed whole genome sequencing on our hiPSC cell line to define a set of 157,552 SNPs occurring within SANCM scATAC-seq peaks for which our SANCMs are heterozygous. We then re-mapped scATAC-seq reads at these loci to reference or alternative alleles and compared the difference in peak accessibility between the two alleles, using a binomial test with correction for multiple hypotheses. Using a cutoff of Benjamini-Hochberg FDR < 0.1, we identified 685 SNPs in 558 candidate regulatory elements that demonstrated chromatin accessibility allelic imbalance (CAAI) in some but not all scATAC-seq clusters, underlying the potential for cell subtype-specific effects by SNPs (Fig. 7a; Supplementary Table 12).

**Figure 7.**
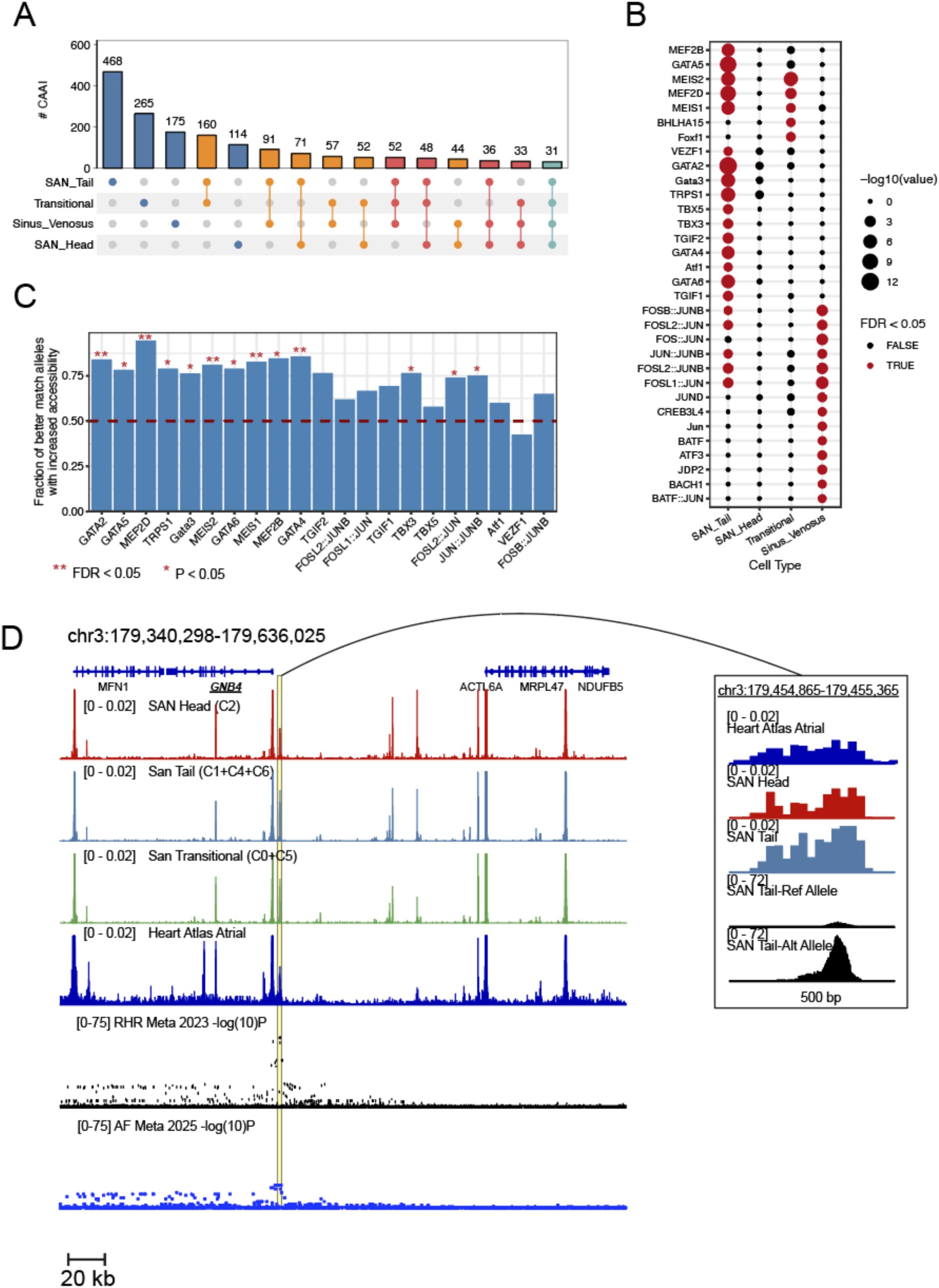
Chromatin Accessibility Allelic Imbalance. (A) The number of SNPs demonstrating chromatin accessibility allelic imbalance (CAAI) in different combinations of annotated cell types. (B) Motif enrichment analysis for transcription factor binding sites predicted to be lost or gained in CAAI sites demonstrates cell type-specific patterns. (C) The percentage of CAAI sites in which the better match for transcription factor binding motifs exhibited increased accessibility. Corrected P values are shown for Fisher’s exact test (‘*’ denotes P < 0.05 and ‘**’ denoted P < 0.01). (D) Chromatin accessibility tracks at the *GNB4* locus from SANCM scATAC-seq re-bulked for SAN head (C2) and SAN tail (C1+C4+C6), SAN Transitional Cells (C0+C5), Heart Cell Atlas atrial cardiomyocytes, with summary statistics from 2 genome wide association study meta-analyses for resting heart rate (RHR) and atrial fibrillation (AF). The region highlighted in yellow corresponds to the scATAC-seq peak in which CAAI was observed and is magnified in the tracks shown on the right, with the addition of 2 tracks corresponding to the SAN Tail – reference allele and SAN Tail – alternate allele. Nearly all the ATAC-seq signal is derived from the alternate allele. Peak heights were normalized and scaled by the ReadsInTSS parameter, the default normalization method in ArchR

Next, we scanned a 40-bp window surrounding each SNP within an accessible chromatin region (including those without CAAI) to define transcription factor binding motifs that were gained or lost in one of the SNP alleles. We then performed motif enrichment analysis to identify motifs more likely to be affected by SNPs exhibiting CAAI than SNPs not exhibiting CAAI (Fig. 7b). In SAN tail cells, we found that MEF2, GATA, MEIS, and TBX3/5 were the most significantly enriched motifs, while in sinus venosus cells, AP-1 motifs were more significantly enriched for CAAI. This agrees with the prevalence of these motifs driving expression in their respective SAN subtypes and underscores the variability of CAAI effects in different subtypes. Next, we asked, for sites in which a transcription factor binding site (TFBS) was lost or gained with variant substitution, whether the allele with the better match to the transcription factor binding motif was associated with greater chromatin accessibility (Fig. 7c). As expected, sites that were better matched with GATA, MEIS, MEF2, TBX, and AP-1 all had significantly *increased* chromatin accessibility, reinforcing a mechanistic model linking common variants in cis-regulatory elements to gene expression: variants that increase affinity for a transcription factor lead to greater chromatin accessibility, increased enhancer activity, and are predicted to lead to higher target gene expression. These data support our hypothesis that CAAI sites are enriched for disease-causing SNPs and support further mechanistic analysis of these sites to better understand disease emergence or progression.

### A Common Variant at the GNB4 Locus Links Heritable Autonomic Sensitivity to Heart Rate and Atrial Fibrillation Susceptibility

The alternate (T) allele of rs7612445(G/T), a SNP exhibiting strong CAAI in SANCMs, is associated with slower resting HR and an increased risk of AF. *Rs7612445* lies within a scATAC-seq peak located 4 kb upstream of GNB4 and is an eQTL for that gene in atrial myocardium with the TT allele exhibiting higher GNB4 expression than the GT and GG alleles.^53^ The chromatin region containing this SNP exhibited the highest allelic imbalance in ATAC-seq reads in SAN Tail, with 90% of reads arising from the alternate allele (Fig. 7d).

To test whether CAAI at this locus translated into a difference in regulatory activity, we cloned the reference and alternate allele into a luciferase reporter vector and transfected both into SANCMs. We found that the construct containing the reference allele repressed promoter activity as compared to the empty vector, whereas the alternate allele, differing only by a single nucleotide, abolished this repressive effect, providing experimental evidence that *rs7612445* regulates chromatin accessibility and transcription near the GNB4 locus (Fig. 8a). To test whether *rs7612445* might alter binding of a critical transcription factor, we identified TFBS predicted to overlap *rs7612445* that are either disrupted or gained by substitution of the alternate allele (Fig. 8b). This analysis predicted a *de novo* MEIS1 site or T-box half-site while simultaneously disrupting a half-site for NR2F1/2, all of which are key regulators in atrial myocardium and in the cardiac conduction system (Fig. 8c). ^54–56^

**Figure 8:**
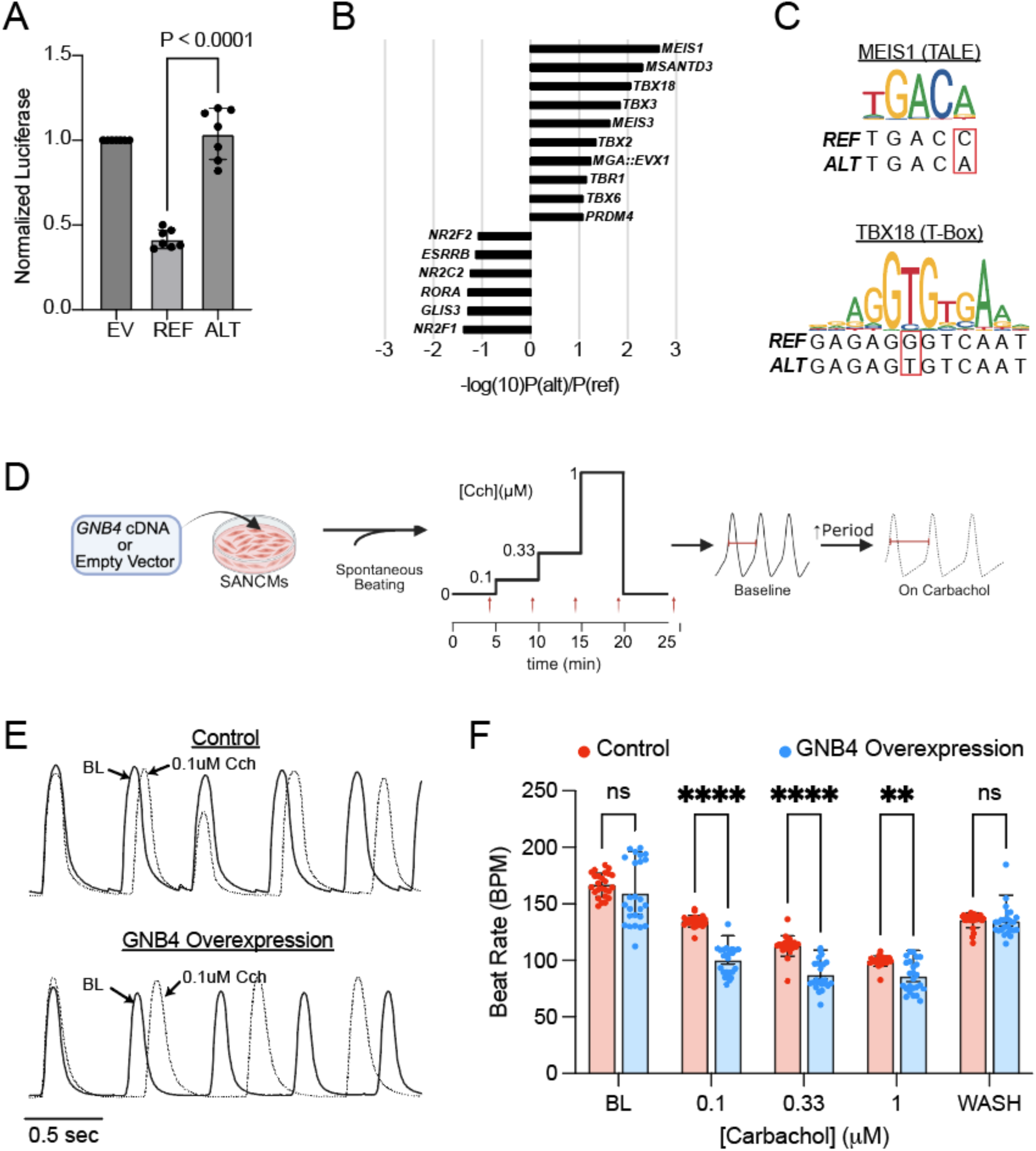
GNB4 Regulates SAN and Atrial Autonomic Response. (A) Luciferase/Renilla comparing the reference and alternative alleles for *rs7612445*. Whereas the reference allele suppresses basal promoter activity, the alternate allele relieves this repression resulting in reconstitution of basal promoter activity (n = 3 separate transfections on 7 distinct differentiations). P values were derived from unpaired T-tests between reference and alternate allele. (B) Ranking of transcription factor binding sites gained and lost with substitution of the alternate allele, defined as those with a greater than 10-fold reduction in the P value (gained site) or 10-fold increase in the P-value (lost site) for a sequence-to-motif match. (C) Motif logo representing the position weight matrices for MEIS1 (JASPAR MA0498.2, above) and TBX18 (JASPAR MA0807.1, below), with the REF and ALT sequences highlighted (red box). (D) Experimental flow diagram (Biorender) representing an assessment of the responsiveness of the beating period of SANCMs to escalating doses of carbachol after transfection of an empty expression vector or an expression vector with *GNB4* cDNA. (E) Raw tracings from the IonOptix imaging system at baseline and in the presence of 0.1 μM carbachol for control cells (transfected with empty vector) and GNB4 overexpressing cells demonstrating cellular edge displacement as a function of time during spontaneous beating, a quantity that was used to estimate the beating rate in the region imaged (F) Spontaneous beating rate of SANCMs with transfected with empty vector (red, control) or a GNB expression vector (blue, GNB4 overexpression) measured at different pixels in different culture dishes. There was no significant difference at baseline or after the wash, but at each carbachol dose assessed, the GNB4 overexpressing cells beat significantly more slowly than the control cells.

As a G-protein beta-subunit, GNB4 is a mediator of cholinergic signaling in the heart through muscarinic acetylcholine receptors. This signaling pathway results in activation of acetylcholine-sensitive potassium channels (IKACh), leading to slower firing rate of PCs in the SAN as well as pro-fibrillatory shortening of atrial action potential duration. We therefore reasoned that the increase in GNB4 expression associated with the alternate allele of *rs7612445* might confer increased sensitivity of SANCMs to cholinergic (parasympathetic) input. To test this hypothesis, we overexpressed GNB4 and measured the spontaneous beating rate of SANCMs after exposure to escalating doses of carbachol (Fig. 8d). Confirming our hypothesis, GNB4-transfected SANCMs exhibited a greater reduction in firing rate as compared to control SANCMs at every concentration of carbachol tested, providing a functional readout of SAN function that plausibly links the alternate allele of *rs7612445* to heritable variation in resting heart rate through its effect on chromatin accessibility at a regulatory element for GNB4 (Fig. 8e,f). Together with the eQTL data and the established association with AF, these data support a model of horizontal pleiotropy, whereby a single common variant regulates distinct heart rhythm-related traits by affecting the sensitivity of both the SAN and the atrial myocardium to parasympathetic input.

## DISCUSSION

In this work, we established that hiPSC-derived SANCMs recapitulate the cellular diversity observed in the human adult SAN and used this system to interrogate heritable variation in regulation of heart rate and susceptibility to atrial fibrillation. Our findings implicate cardiac inherited autonomic responsiveness as a mechanism that connects sinus node function to arrhythmia susceptibility, providing a clear example of the utility of this approach to elucidate mechanistic relationships among closely related cardiovascular traits.

### Establishing a Model System for SAN Cardiomyocytes

To generate SANCMs, we used growth factors and morphogens to recapitulate signaling pathways involved in SAN development, including biphasic Wnt signaling, retinoic acid, and inhibition of TGFβ and FGF signaling^24,34^, yielding a cell population that compared favorably with other approaches, as evidenced by gene expression and electrophysiological behavior. Subclustering of these SANCMs defined DEGs that mark each PC subtype. These DEGs significantly overlapped with an scRNA-seq dataset derived from mouse neonatal SAN^57^, indicating conservation of the gene regulatory networks controlling compartmentalization of the SAN. While previous work demonstrated that directed differentiation recapitulated the developmental trajectories adopted by differentiating embryonic venous pole cardiomyocytes ^27^, we extend these important findings by showing that single cell expression profiles and epigenetic profiles from SAN subtypes recapitulate expression of key marker genes and noncoding regulatory elements in the adult human heart. Bidirectional mapping of SANCMs to the human cell atlas showed strong concordance at the level of cellular subtypes, indicating that essential features of the gene expression programs driving specialization of cardiomyocytes are present in relatively immature hiPSC-SAN cultures.^38^ This finding is important because many common variants associated with traits ascertained in adulthood (such as resting heart rate and atrial fibrillation) are likely to regulate gene expression in the adult heart.

### Transcriptional Regulators in SAN Cell Subtypes

Aggregating scATAC-seq data by cluster demonstrated differentially accessible chromatin regions associated with different PC subtypes and identified established enhancers that were previously known for SAN transcription factors SHOX2, ISL1, and TBX3, as well a novel enhancer for TBX18 that is more active in SAN head^36,39^. By demonstrating differential enhancer accessibility between SAN head, SAN tail, and transitional cells, our data suggest that the signaling upstream of these established SAN enhancers is involved in both specification as well as the maintenance of fine nodal patterning. In addition, motif enrichment analysis highlighted the significance of AP-1 motifs in peaks from sinus venosus myocardium, suggesting that this pathway could be critical for differentiation of this cell type from SAN myocardium. Of note, sinus venosus myocardium exhibited a hybrid cardiomyocyte-smooth muscle gene program, so one possibility is that AP-1 may promote formation of TBX18+ progenitors and that persistent AP-1 activity could direct TBX18+ progenitors towards a sinus venosus fate. ATAC-seq data from mouse SAN revealed that AP-1 motifs are highly overrepresented in differentially accessible chromatin in neonatal mouse PCs as compared to atrial myocardium, suggesting an evolutionarily conserved role for AP-1^36^.

The NFI motif was also the most highly overrepresented motif in SAN head compared to tail myocardium and both *NFIA* and *NFIB* expression were increased in SAN head cells. Further supporting a role for this family of TFs in SAN subtype differentiation, comparison of our scRNA-seq with a mouse dataset demonstrated that *NFIA* is differentially expressed in mouse SAN head versus SAN tail cells, a finding we confirmed by establishing co-expression of NFIA and NFIB in the murine developing SAN head^57^. Although a recent study showed that loss of NFIX transcription factor in mice results in altered adult SAN physiology without other effects on cardiac structure and function^58^, NFI transcription factors have not been previously explored in SAN development and patterning. With ISL1, TBX, and MEF2, our motif enrichment analysis supports a role for NFI as an activating transcriptional regulator in the SAN Head requiring further exploration.

### HiPSC SAN Cultures for Functional Genomics

Common variants are theorized to influence traits by directly affecting the activities of *cis*-regulatory elements that regulate expression of nearby genes within functionally relevant cell types^59,60^. Consistent with this framework, we found that differentially accessible chromatin in each of the SANCM subtypes significantly overlapped with SNPs associated with cardiac traits such as resting heart rate (SAN tail and transitional cells > SAN head) and heart rate recovery after exercise (SAN head > SAN tail). This finding is supports the model of functional compartmentalization, in which migration of the leading pacemaker along the cranial-caudal dimension is driven by changes in autonomic tone that affects SAN head cells disproportionately, whereas resting heart rate is driven by mechanisms of impulse generation and transmission that depend on function of the SAN tail and transitional cells.

One of the strongest trait enrichments we observed was for AF. Clinically, there is a well- established link between sinus node function and the propensity to develop AF^61^. Because of the existence of common risk factors for both conditions, determining the causal basis for this association has been a long-standing question in the field with major clinical implications. More recent evidence indicates that heart rate determination and AF may have overlapping genetic architectures^19^, although the ability to explore the mechanistic basis for these findings has been limited. Supporting this notion, we found enrichment of OSR1 binding motifs in AF-associated candidate regulatory elements in SANCMs. During cardiogenesis, *OSR1* is expressed transiently in the posterior second heart field where it is required for morphogenesis of the cardiac venous pole, interatrial septum, and interface between the SAN and the atrium ^46–48^. Enrichment of OSR1 motifs in AF-associated accessible chromatin regions from relevant cell subtypes provides evidence for a developmental contribution to the shared genetic architecture of AF and SAN function.

Previous studies have leveraged expression (eQTL analysis), chromatin accessibility, allele specific affects in ChIP-Seq, and annotation of functional regulatory elements (e.g. ENCODE) to map SNPs that are more likely to be causative ^62 63–65^. In addition, functional genomics of atrial fibrillation^66^ and QT interval^67^ have been explored in murine HL-1 cardiomyocytes as well as congenital heart disease loci ^68^ in hiPSC-derived cardiomyocytes using lentiviral parallel reporter assays. These studies have identified conserved gene regulatory elements that control expression of rhythm-related genes and have detected allele-specific effects on enhancer activity.

Here we leveraged an *in vitro* system in which rare human cell subtypes are represented and are thus more amenable to human genomic studies, reporter assays and functional assays including physiological processes. Although hiPSC-derived cardiomyocytes are immature phenotypically, the concordance of gene expression and accessible chromatin regions with adult human counterparts from the Cell Atlas project motivated our mechanistic exploration of heart rhythm-related genetic variation in our hiPSC SANCMs. Specifically, we tested whether we could identify novel functional regulatory elements at genomic loci associated with heart rhythm, and whether the system is sensitive enough to detect effects of individual common variants and to distinguish them from variants merely associated with traits through linkage to causative variants.

### snpSTARR-Seq in SANCMs

Refinements of STARR-seq have been used to quantify enhancer strength among smaller subsets of putative regulatory elements (capSTARR-seq)^49^ and to discern the effects of single variants on enhancer activity (snpSTARR-seq) ^50^. Indeed, STARR-seq was used in a previous study to identify regulatory elements and allelic effects at several atrial fibrillation-associated loci using a rat atrial cell line (near genes PITX2, PRRX1, SCN5A/10A, WNT8A, GJA1, CAV1, NEURL, SYNE2, HCN4, and ZFHX3) ^69^. Here we have advanced this assay by applying it to a human iPSC-derived culture that contains several cellular subtypes exhibiting transcriptomic and epigenetic fidelity to their adult *in vivo* counterparts. This approach allowed us to broaden the range of potentially significant loci in cell types that may be directly relevant to human heart rhythm and AF.

Most of the candidate regulatory elements we examined displayed self-transcribing activity in SANCM cultures when compared to negative control regions, while a small percentage showed effects of individual alleles on enhancer activity in STARR-seq, a finding in agreement with the notion that only a fraction of lead SNPs are functional and causal for traits. The excellent reproducibility of the assay over 5 replicates with low variance and the ability to detect allelic imbalance would lend support to a more general effort to include all ATAC-peaks in the genome containing a heart rhythm-associated SNP, possibly coupled with single cell sequencing to ascertain cell type specific enhancer activity and cell type specific allelic effects.

In addition, a SNP within an enhancer at the *ATXN1* locus exhibits allelic imbalance in self-transcription. ATXN1 is a transcriptional co-repressor ^70^ without a well-established function in the heart, although loss of function in mice leads to deficits in neurological ^71^ and immune system function ^72^, and trinucleotide repeat expansion in *ATXN1* is the cause of spinocerebellar ataxia type I ^73^. The alternate allele of this SNP creates a nuclear receptor consensus site and improves the local affinity for bZIP family transcription factors, leading to an increase in constitutive enhancer activity in hiPSC-SANCMs as well as signal responsiveness. In theory the increase in basal activity could be conferred by a factor binding to the nuclear receptor site such as NR2F2, a regulator of atrial gene expression ^54^, while CREB1/ATF/AP-1 and NR4A, regulators of cardiac stress response^74^, could contribute to the signal responsiveness. While further work will be required to identify the specific proteins that bind this enhancer in atrial myocardium, our findings support the hypothesis that ATXN1 expression in cardiomyocytes may protect against atrial fibrillation and provide a physiological context for exploring the gene-trait relationship further. In addition, this example illustrates the potential for functional genomics to provide mechanistic information beyond mere association.

### Use of Chromatin Accessibility Allelic Imbalance for Fine Mapping

CAAI offers a complementary approach to identify sites genome-wide where variants are associated with chromatin accessibility. SNPs associated with CAAI are theorized to be enriched for causative variants that drive the difference in chromatin accessibility by altering transcription factor binding affinity. While allelic affects have been observed previously using genome wide epigenetic binding data, coupling CAAI with scATAC-seq allows for deeper biological insight by distinguishing the specific cell subtypes that are most relevant to the association between the CAAI-associated SNP and the trait ^75^. Indeed, only a minority of SNPs exhibiting CAAI in our analysis did so in all cell types we examined. Most CAAI sites were restricted to one or a few cell subtypes, underscoring the highly contextual nature of transcription factor binding to sites driving genetic variation. SAN Tail cells had the most sites with CAAI which may reflect that their transcription program merges those of both atrial cardiomyocytes and SAN head cardiomyocytes and thus engages a broader set of enhancers. Further supporting the value of coupling CAAI with single cell epigenetic data, cell type-specific transcriptional signatures were observed at CAAI sites predicted to disrupt transcription factor binding sites. These findings reinforce the significance of the cell type specific transcriptional hierarchies not only in driving cellular specialization but in mediating heritability of traits. We also found that in CAAI sites with predicted TFBS disruptions, the allele with the better match to the TFBS more commonly exhibited greater accessibility. This result supports the interpretation that many of the CAAI SNPs represent functional regulatory variants. Finally, while CAAI was effective for fine mapping trait-associated loci identified with GWAS, its use was limited to SNPs for which our iPSC line was heterozygous. In theory, this limitation could be overcome by repeating our analysis using hiPSCs-SANCMs derived from a larger group of individuals representing more haplotypes. Broader application of this powerful tool could facilitate genome-wide filtering of SNPs to those most likely to be causal while suggesting specific biological mechanisms through examination of disrupted TFBS. Finally, differentiating SANCMs from male and female subjects could shed light on established sex differences in heart rate regulation and propensity for tachycardia, a clinically relevant but poorly understood area ^76^.

### Role of Autonomic Sensitivity in Mediating the Association of Sinus Node Function and Atrial Fibrillation

Genes required for mediating autonomic responsiveness of the SAN are highly enriched among GWAS hits for heart rate determination and heart rate response to exercise ^77^. Not surprisingly, we found that scATAC-seq peaks at many of these loci contained SNPs associated with heart rate. Furthermore, the association of scATAC-seq peaks from all myocardial cell types with atrial fibrillation risk suggested that the SANCM platform might offer a means to explore the genetic contribution to the association between sinus node function and atrial fibrillation.

It is well established that vagal input to the atrium and pulmonary venous myocardium can facilitate atrial fibrillation through IKACh, an outward hyperpolarizing current that is activated by Gβγ in response to muscarinic agonism by acetyl choline or local adenosine release ^78,79^ . In atrial myocardium, IKACh activation leads to action potential shortening and thereby to a shorter refractory period and an improved safety factor for conduction, both of which are pro-fibrillatory changes in tissue electrophysiology. Indeed, the drug Tertiapin, a specific IKACh blocker, is effective for termination of atrial fibrillation episodes in some contexts ^80^. In the SAN, the same pathway is operative, but the hyperpolarization caused by IKACh leads to a reduction in the slope of diastolic depolarization and a lower maximum diastolic potential, both of which reduce automaticity ^81,82^. Thus, a heritable increase in cardiac vagal sensitivity would, in theory, result in a higher risk of atrial fibrillation and a lower heart rate.

Our finding of CAAI in a regulatory element upstream of GNB4 provides an illustrative example of this phenomenon. When tested in a luciferase assay, the regulatory element containing the alternate allele of this enhancer increased transcription in comparison to the reference allele. Thus, our data would predict that the alternate allele should result in more GNB4 expression, a finding that has been confirmed in a previous work ^53^. GNB4 encodes for a G protein beta subunit expressed in the heart. Using our hiPSC system, we therefore defined functional consequences of increased GNB4 for autonomic sensitivity of SANCMs and we observed exaggerated heart rate slowing induced by a muscarinic agonist in cells overexpressing GNB4. Taken together, these findings suggest a mechanism of horizontal pleiotropy, whereby a single SNP influences distinct physiological traits in the atrium and SAN to drive a genetic association between these two traits, a result that would be difficult to discern in the absence of a cell type resolved functional genomics platform.

## ONLINE METHODS

### Cell line and maintenance of hiPSC

All differentiations were performed using a commercially available Human episomal iPSC line (ThermoFisher). Culture of hiPSCs was carried out on hESC-Qualified Matrigel (Corning) coated dishes with mTeSR Plus medium (STEMCELL Technologies) and was refreshed every day. Cells were passaged at 80-90% confluence (∼every 4 days) using ReLeSR (STEMCELL Technologies) according to manufacturer’s instructions.

### Differentiation of hiPSCs

Monolayer differentiations of cardiomyocytes were carried out using a modified GiWi strategy (Lian et al 2013). hiPSCs were grown on Matrigel to 90% confluency then harvested as single cell suspension using TrypLE (ThermoFisher) and resuspended in mTeSR Plus containing 10µM ROCK inhibitor Y-27632 (Reprocell). Cells were counted using a Vi-CELL XR Cell Viability Analyzer (Beckman Coulter) and re-plated onto Matrigel coated plates at 15,625-18,230 cells/cm^2^ in mTeSR Plus containing 10µM ROCK inhibitor Y-27632 (day -3 of differentiation). Cell plating density was re-optimized every 10 passages or upon a new thaw. At day -2 media was changed to mTeSR Plus. For ventricular cardiomyocytes, at day 0 when cells were 80-90% confluent media was changed to Cardiac Differentiation Media (CDM; RPMI1640 with 25mM HEPES and Glutamax (ThermoFisher) and 1x B27 minus insulin (ThermoFisher)) containing 8µM CHIR99021 (Sigma). On day 2 of differentiation cells were changed to CDM containing 5μM IWP2 (Sigma). At day 4, media was changed to CDM until day 6 when media was switched to Cardiac Maintenance Media (CMM; RPMI1640 with 25mM HEPES and Glutamax (ThermoFisher) and 1x B27 supplement (ThermoFisher). Media was changed every 2 days until day 20. On day 20 cells were split 1:4 in preparation for metabolic selection. Cells were washed with TrypLE for 5 minutes at room temperature then digested into single cell suspension using prewarmed Accumax (Sigma) for 20-30 minutes at 37°C. Digestion was quenched using CMM containing 20% Knock-out serum replacement (ThermoFisher) and 10µM ROCK inhibitor Y-27632 and cells collected by centrifugation at 200xg for 5 minutes. Cells were then resuspended in CMM containing 20% Knock-out serum replacement and 10µM ROCK inhibitor Y-27632 and plated onto new Matrigel coated dishes at 25% the original confluence (1:4 split). 24 hours later (day 21) media was changed to CMM. At day 23 media was changed to Cardiomyocyte Selection Media (CSM; RPMI1640 with Glutamine and No glucose (ThermoFisher), 5mM Sodium DL-lactate solution (Sigma) and 0.5mg/ml Human Serum Albumin (Sigma)). CSM was changed every 2 days until day 31 when cells were switched back to CMM. Cells were maintained in CMM (media changed every 2 days) until cells were either frozen (day 34) or used for analysis.

For atrial cardiomyocytes, the same protocol was followed except for the following addition. On day 3 of differentiation, 1µM Retinoic Acid (RA; Sigma) was spiked into the CDM containing IWP2. For SAN cardiomyocytes the ventricular protocol was following with the following modifications. At day 3, 2.5ng/ml BMP4 (R&D Systems), 5.4µM SB431542 (Tocris) and 0.5µM RA were spiked into the CDM containing IWP2. On day 4, media was replaced with CDM containing 240nM PD173074 (Tocris).

Cryo-preserved cardiomyocytes were recovered in CMM containing 20% Knock-out serum replacement (ThermoFisher) and 10 µM ROCK inhibitor Y-27632 and plated on Fibronectin (5 µg/cm^2^, for TC-treated plates or 10 µg/cm^2^, for glass bottom plates) in Gelatin (0.1%) at a density of 200-250k cells/cm^2^. The next day media was changed to CMM and exchanged every 48 hours thereafter. Cells were allowed to recover for at least 3 days prior to use in any assay.

### Quantitative real-time RT-PCR

Total RNA was extracted from cells using RNeasy mini kit (Qiagen) according to the manufacturer’s instructions. Purified RNA was subjected to reverse transcription as described in the SuperScript IV VILO with ezDNase enzyme kit (ThermoFisher). Following cDNA synthesis, expression level of the genes was quantified by quantitative Real-Time PCR (qPCR) using TaqMan Fast Advanced Master Mix (ThermoFisher) on a QuantStudio 7 Flex Real-Time PCR system (ThermoFisher). The quantification was analyzed using the C_T_ (threshold cycle) values. Target gene expression was normalized to *GAPDH.* Expression was then calculated using the 2^−ΔΔCt^ method and graphed relative to ventricular cardiomyocytes.

### Immunofluorescence staining

Cells were cultured in CellCarrier-96 Ultra Microplates (Perkin Elmer) coated with 50µg/ml fibronectin (Sigma) and 0.1% Gelatin (STEMCELL Technologies). Cells were washed with PBS then fixed with 4% PFA for 20 min at room temperature and washed again with PBS. Cells were then blocked and permeabilized with 0.1% Triton X-100 and 5% donkey serum in PBS for 1 hour at room temperature on a rotator. Primary antibodies diluted in blocking/permeabilization buffer were added and incubated at 4°C overnight on a shaker at 3000 rpm. Cells were washed 3 times for 3 minutes in PBS at room temperature on a shaker at 3000 rpm. Secondary antibodies diluted in PBS with 5% donkey serum were added and incubated at room temperature for 1 hour shaking at 3000 rpm then followed by 2×3 minute washes in PBS with shaking. The cell nuclei were counterstained with Hoesht 33342 (1:2000; Thermo Fisher) for 10 minutes with shaking and then washed 2×3 minutes with shaking in PBS. Cells were imaged with an Opera Phenix High Content Screening System (Perkin Elmer) using Harmony software and analyzed using Columbus software (Perkin Elmer).

The following primary antibodies were used: mouse anti-Cardiac Troponin T (ThermoFisher; 1:200), rabbit anti-Cardiac Troponin T (Abcam; 1:200), mouse anti-ISL1/2 (DSHB; 1:200), rabbit anti-NKX2.5 (Cell Signaling; 1:200), rabbit anti-HCN4 (Alomone; 1:200) and mouse anti-SHOX2 (Abcam; 1:200). The following secondary antibodies were used: donkey anti-mouse Alexa Fluor 488 (ThermoFisher; 2µg/ml) and donkey anti-rabbit Alexa Fluor 555 (ThermoFisher; 2µg/ml).

### Imaging of Voltage-Sensitive Dye

Voltage imaging of VCMs and SANCMs was performed by adding FluoVolt membrane potential dye (ThermoFisher) to 96 well-plates seeded with 100k cells/well. Briefly, FluoVolt was mixed 1:20 with PowerLoad concentrate, and diluted 100x with FluoroBrite DMEM (ThermoFisher). 100x water-soluble Probenecid (ThermoFisher) was then added to achieve a final concentration of 1x. After washing with FluoroBrite DMEM, cells were incubated for 5 min at RT with the dye solution. The dye was then removed and replaced with FluoroBrite DMEM with 1x Probenecid. The cell plate was then placed inside the culture chamber of an iMX4 microscope (Molecular Devices) and equilibrated for 5 minutes. Time lapse images were then acquired for 5 seconds per site at 100 fps, using a 10x objective. For Zatebradine measurements, cells were prepared as described above then treated with 0, 0.1, 0.3, 1 or 3 µM Zatebradine (Tocris), incubated for 5 minutes then imaged. The images (5 fields per well, 4 wells per treatment) were then analyzed using a custom Matlab script that extracted individual voltage traces for each cell in the field and calculated electrophysiological parameters including beating rate, amplitude, and action potential duration at 20%, 50% and 90% repolarization. All parameters were averaged and compared between groups using a student’s T-test after each distribution passed a normality test.

### scRNA-seq data analysis

Initial processing of scRNA-seq data was done with the Cell Ranger Pipeline (https://support.10xgenomics.com/single-cell-geneexpression/software/pipelines/latest/what-is-cell-ranger, v.6.0.0) by first running cellranger mkfastq to demultiplex the bcl files and then running cellranger count with 10x Genomics’ pre-built Cell Ranger reference GRCh38-2020-A_build. After running Cell Ranger, the filtered_feature_bc_matrix produced by Cell Ranger was read into R (v.4.0.4) with the Seurat (v.4.3.0) function Read10X ^83^. scDblFinder (v.1.2.0) was run for each sample for detecting doublets ^84^. Cells classified as doublets were not enriched in any clusters identified in the downstream clustering analysis and thus retained. Cells from all samples were merged into a single Seurat object and analyzed separately for each dataset.

For the analysis of cells of day 34 datasets (ACM, VCM, and SANCM cultures), cell barcodes were filtered based on the number of genes per cell (between 1,000 and 9,000), unique molecular identifiers (UMIs) per cell (more than 1500), and percentage of mitochondrial reads per cell (less than 50%). The filtered cell number used for subsequent analysis was 20,712 cells for the day 34 SANCM culture, 21,676 cells for the day 34 ACM culture, 21,934 cells for the day 34 VCM culture, 102,496 cells for the SANCM time-course day 3 to day 34 cultures, respectively.

Principal component analysis (PCA) was performed using the RunPCA function and the top 40 principal components (PCs) were selected. UMAP was computed using the RunUMAP function and cells were clustered using a k-nearest neighbor graph and the Louvain algorithm with the FindNeighbors and FindClusters functions. A resolution of 0.6 and 1.0 was used for day 34 datasets and time-course datasets, respectively. Expression of canonical marker genes and enriched in the marker genes (described below) in the resulting clusters were then used to label clusters.

For the sub-clustering analysis of day 34 SANCM, ACM, and VCM cultures, replicates were integrated using the Seurat integration procedure. When running FindIntegrationAnchors and IntegrateData, the normalization.method parameter was set to the “SCT” using 30 dimensions and 5000 features. For dimensionality reduction, the integrated expression matrix was used to compute PCs by RunPCA. The top 40 PCs were selected for UMAP visualizations and clustering analysis. The cells were then clustered using Seurat’s FindNeighbors and FindClusters with a resolution of 0.5 for the day 34 SANCM dataset, 0.4 for the day 34 VCM dataset, and 0.3 for the day 34 ACM dataset, respectively. The clusters were manually annotated based on canonical markers and enriched pathways.

For the sub-clustering analysis of the day 34 SANCM dataset, cells were first separated into 20 clusters as described above. Eleven clusters were removed due to enrichment of cell cycle genes (3 clusters), low gene count (4 clusters), high percentage of mitochondrial reads (2 clusters), enrichment of genes in cell stress related pathways (1 cluster), or without consistent expression of canonical markers of any cell type (3 clusters). Cells in the retained clusters were normalized and integrated again using the same pipeline. Dimensionality reduction was recomputed, and clustering analysis was performed with a resolution of 0.8.

For the sub-clustering analysis of SAN head and tail cells, the three clusters identified as SAN head and tail cells (cluster 2, 0, and 3 in Fig. 2a) were subset and analyzed separately. Normalization was performed by Seurat’s sctransform method with the 3,000 most variable genes and regressing out cell cycle effects, and batch effect was removed using Harmony (v. 0.1.0)^85^.Cluster identification and UMAP reduction were conducted as above, except that a resolution of 0.1 was used for the clustering.

For each dataset, the FindAllMarkers and FindMarkers functions were performed after normalizing the expression counts with Seurat’s “LogNormalize” method to identify the marker genes for each cluster (min.pct = 0.25, logfc.threshold = 0.25) and intercluster differentially expressed genes (min.pct = 0.25, Padj < 0.05, logfc.threshold = 0.5 for SANCM time-course datasets and 0.25 for day 34 datasets) via the Wilcoxon rank sum test. Pathway analysis of marker genes and differentially expressed genes was conducted with MetaScape (v3.5) ^86^.

### Integration of publicly available scRNA-seq datasets

One preprocessed scRNA-seq dataset was downloaded from the Gene Expression Omnibus (GEO) (accession code GSE189782^27^) and processed using Seurat as described in the studies. Processed snRNA-seq data of human SAN from donors with normal conduction parameters^38^ was downloaded from the Heart Cell Atlas (https://www.heartcellatlas.org). We day performed reference mapping to project the cells in the two public datasets onto the UMAP structure of our day 34 SANCM scRNA-seq data according to Seurat’s “Mapping and annotating query datasets” tutorial (https://satijalab.org/seurat/articles/integration_mapping.html). Specifically, the FindTransferAnchors function was used to find a set of anchors between the reference and query object with the top 40 PCs and “SCT” normalization. These anchors were then passed into the MapQuery function to project the query data onto the UMAP embedding of the reference.

Processed Visium spatial transcriptomics data of OCT-embedded human SAN samples from 8 donors with normal conduction parameters was downloaded from the Heart Cell Atlas (https://cellgeni.cog.sanger.ac.uk/heartcellatlas/v2/visium-OCT_SAN_raw.h5ad). ^38^ To spatially map day 34 SANCMs in these Visium data, we used cell2location (v.0.1.3), ^87^ a Bayesian model that decomposes spatially resolved multi-cell RNA counts matrices into the reference signatures, thereby establishing a spatial mapping of cell types. Briefly, we first estimated reference signatures of cell types using our day 34 SANCM scRNA-seq data with cell2location’s negative binomial regression model. The inferred reference signatures were used for cell2location to decompose mRNA counts in each Visium spot into the abundance of each reference cell type. The single-cell negative binomial regression was trained with default settings after selecting genes with parameters nonz_mean_cutoff = 1.12, cell_count_cutoff = 5, and cell_percentage_cutoff2 = 0.03. The deconvolution model was trained with parameters max_epochs = 30,000, detection_alpha = 20, and n = 5.

### scATAC-seq data analysis

Initial processing of the day 34 SANCM scATAC-seq data was performed using Cell Ranger ATAC (https://support.10xgenomics.com/single-cell-atac/software/pipelines/latest/what-is-cell-ranger-atac, v.2.0.0) by first running cellranger-atac mkfastq to demultiplex the bcl files into FASTQ files and then running cellranger-atac count to generate scATAC fragments files with 10x Genomics’ pre-built Cell Ranger reference GRCh38-2020-A_build and default parameters.

These fragments files were loaded into R (v.4.0.4) using the createArrowFiles function in ArchR (v.1.0.1)^88^. Quality control metrics were computed for each cell, and cells with TSS enrichments less than 4 and number of unique fragments less than 1,000 were filtered out for all samples. The number of cells retained was 38,744 (ranging from 10,565 to 15,823). Doublets were detected by AMULET (v.1.1) ^89^ for each sample. Cells classified as doublets were retained due to lack of enrichment in any clusters identified in the subsequent clustering analysis.

After generating arrow files for each sample, an ArchR project containing all samples was created. As part of construction of the arrow files, a tile matrix consisting of reads in 500-bp tiles was constructed. This tile matrix was used as input to compute an iterative latent semantic indexing (LSI) dimensionality reduction using the addIterativeLSI function in ArchR with a total of 2 iterations, a clustering resolution of 0.2 following the first iteration, 50,000 variable features, 30 dimensions and sampling 10,000 cells. Harmony (v. 0.1.0)^85^ was applied to the dataset to correct for the batch effects. Using the Harmony-corrected dimension reduction, clustering was performed using ArchR’s addClusters with a resolution of 0.45. We next ran addUMAP on the Harmony-corrected dimensionality reduction with 30 nearest neighbors and a minimum distance of 0.5. Gene activity scores computed by ArchR based on the accessibility within and around a given gene body were examined for the known marker genes. Clusters with less than 34 cells consisted mainly of cells from only a subset of the samples and were removed from the subsequent analysis. One cluster containing about 3000 cells that could not be confidently assigned to one cell type and probably constitute poor quality nuclei was also removed from further analysis.

For sub-clustering of SAN head and tail cells, scATAC-seq clusters identified as SAN head and tail cells were subset and used to recompute iterative LSI dimensionality reduction as described above. Batch correction, cluster identification, and UMAP reduction were also performed as above, except that a resolution of 0.2 was used for the clustering.

### Integration of scATAC-seq and scRNA-seq data

day 34 SANCM scRNA-seq gene expression counts were integrated with scATAC-seq data with addGeneIntegrationMatrix in ArchR as a constrained integration, grouping clusters together during addGeneIntegrationMatrix to the following groups: SAN head, SAN tail, transitional, Sinus Venosus, proepicardial and neuronal cells. The output of the integration step resulted in scATAC-seq cells having both a chromatin accessibility and a gene expression profile. Accessibility gene scores and transferred RNA expression counts were imputed with addImputeWeights.

### Cell-type-specific peak and TF motif enrichment

Pseudobulk group coverages of cell type clusters were calculated with the ‘addGroupCoverages’ function and used for peak calling using the ‘addReproduciblePeakSet’ function in ArchR. A background peak set controlling for total accessibility and GC content was generated using addBgdPeaks for TF enrichment analyses. Cell type–specific marker peaks and differentially accessible peaks between clusters were identified with ‘getMarkerFeatures’ with the Wilcoxon test (maxCells = 10,000, FDR ≤ 0.05, and Log2FC ≥ 0.5 for marker peaks, and maxCells =6,000, FDR ≤ 0.01, and |Log2FC| ≥ 1 for differential accessible peaks). The enriched motifs for marker peaks of each cluster and differential accessible peaks were predicted using the ‘addMotifAnnotations’ function in ArchR based on the cisbp motif set. Chromvar was run with addDeviationsMatrix to calculate enrichment of chromatin accessibility at different TF motif sequences in single cells. To identify correlations between the gene expression and TF activity, the GeneIntegrationMatrix and the Chromvar deviations (MotifMatrix) were correlated using correlateMatrices. A correlation of greater than 0.5 and an adjusted p-value less than 0.01 were used to determine whether TF expression and activity were positively correlated. “Peak-to-gene links” were calculated using correlations between peak accessibility and integrated scRNA-seq expression data using addPeak2GeneLinks with the parameter ‘maxDist = 1.5e6’ and getPeak2GeneLinks in ArchR. We further visualized the peak to gene links using the ‘plotBrowserTrack’ function in ArchR.

### Integration of GWAS and scATAC-seq data

GWAS results for resting heart rate, heart rate response to recovery post exercise, and atrial fibrillation were used to integrate with scATAC-seq data. First, summary statistics for resting heart rate (GCST007609), and heart rate response to recovery post exercise (GCST005847), and atrial fibrillation (accessions codes GCST006414 ^42^ and GCST006061^43^) were downloaded from the GWAS catalog. All the SNPs were then assigned to their nearest genes based on their coordinates. Bedtools (v.2.30.0)^90^ was used to intersect the SNP coordinates with marker peaks of SAN tail for resting heart rate, SAN head for heart rate response to recovery post exercise, and transitional cells for atrial fibrillation respectively and also the union of all the peaks identified in these SAN cells.

### Stratified linkage disequilibrium (LD) score regression

We used the LDSC package (v1.0.1) ^60^ to estimate genome-wide GWAS enrichment for a variety of phenotypes within the cell type-specific scATAC-seq peaks. We first downloaded GWAS summary statistics for: atrial fibrillation, resting heart rate, heart rate increase in response to exercise, heart rate response to recovery post exercise (20 sec) (the GWAS catalog https://www.ebi.ac.uk/gwas/; accessions codes GCST006414, GCST007609, GCST005845, and GCST005847); height (https://rpubs.com/leaknielsen/1027689); diastolic blood pressure, systolic blood pressure, years of education, smoking status, auto immune traits, asthma, dermatologic diseases, high cholesterol, type 2 diabetes, age at menarche, BMI (https://alkesgroup.broadinstitute.org/UKBB/); schizophrenia, major depression disorder, Alzheimer’s disease (https://pgc.unc.edu/for-researchers/download-results/; pubMedIDlink 25056061, 29700475, 30617256); coronary artery disease, QT prolongation, LV stroke vol. indexed, sscending aorta diameter index, LV max. area index, LV min. area index, pulmonary artery:aorta ratio, RA fractional area change, RA max. area index, RA min. area index, RV end diastolic vol. index, RV ejection fraction, RV end systolic vol. index, P-wave duration, PR interval, QRS duration, QTc interval, RR interval, heart failure, LV ejection fraction (https://personal.broadinstitute.org/ryank/; Aragam_2022_CARDIoGRAM_CAD_GWAS.zip, Nauffal_2022_QT_GWAS_SAIGE.zip, Pirruccello_2022_UKBB_HeartStructures.zip, Choi2021_TOPMed_freeze6.zip, HERMES_Jan2019_HeartFailure_summary_data.txt.zip, MRI_lvef_filtered.zip); IBD (https://www.dropbox.com/s/ttuc6s7tv26voq3/iibdgc-trans-ancestry-filtered-summary-stats.tgz?dl=0); total cholesterol (http://csg.sph.umich.edu/abecasis/public/lipids2013/jointGwasMc_TC.txt.gz). We then used the provided munge_sumstats.py script to convert these GWAS summary statistics to a format compatible with LDSC.

Using all the ATAC-seq peaks called within each cluster, we generated the annotation-specific partitioned LD score files for each cluster separately according to the “Partitioned LD scores” section of the LD score estimation tutorial (https://github.com/bulik/ldsc/wiki/LD-Score-Estimation-Tutorial). We downloaded the list of 1,154,611 candidate cis-regulatory elements (cCREs) for 222 distinct cell types from the Human Enhancer Atlas (http://catlas.org/humanenhancer/#!/)^91^ and used them as background annotations. We lifted over peak coordinates from hg38 to hg19 for both day 34 SANCM ATAC-seq peaks and background peaks using CrossMap (v.0.6.3) ^92^ before computing annotation-specific LD scores. We then used these hg19 bed files to make annotation files for each cell type. For each trait, we used LD score regression to estimate coefficient p value for each cell type relative to the background annotations according to the cell-type-specific analysis tutorial (https://github.com/bulik/ldsc/wiki/Cell-type-specific-analyses) and used the Benjamini-Hochberg correction for multiple testing.

### Luciferase assays

VCMs and SANCMs were recovered and plated in white walled clear-bottom 96-well assay plates (Corning). On the day of transfection media was changed to CMM with 10% FBS. Transfections were carried out using Viafect transfection reagent (Promega) at a ratio of 6:1 (µl Viafect: µg DNA). Transfection complexes were formed in Opti-MEM serum free media (Thermo-Fisher) according to the manufacturer’s instructions using 100 ng pGL4.23 (enhancer-minP-luc2, Promega) and 5 ng pGL4.74 (hRluc, Promega) per well and performed in triplicate. 24 hours post-transfection, media was changed to CMM for an additional 24 hours before performing the luciferase assay. Luciferase measurements were carried out using the Dual-Glo Luciferase Assay System (Promega) following the manufacturer’s protocol and using a Glo-Max Explorer microplate reader. Luminescence values were first normalized to Renilla activity then expression was calculated relative to the empty vector control.

### Reporter transfection and Fluorescence-Activated Cell Sorting (FACS)

SANCMs were recovered and plated in 6-well plates (Corning). On the day of transfection media was changed to CMM with 10% FBS. Transfections were carried out using Viafect transfection reagent (Promega) at a ratio of 6:1 (µl Viafect: µg DNA). Transfection complexes were formed in Opti-MEM serum free media (ThermoFisher) according to the manufacturer’s instructions using 3.2 ug of plasmid containing TBX18-R3 enhancer-minP-GFP per well. 24 hours post-transfection, media was changed to CMM for an additional 24 hours before cells were harvested for FACS. Cells were digested into single cell suspension using prewarmed Accumax (Sigma) for 20-30 minutes at 37°C. Digestion was quenched using CMM containing 20% Knock-out serum replacement (ThermoFisher) and 10 µM ROCK inhibitor Y-27632 and cells collected by centrifugation at 200xg for 5 minutes. Cells were then resuspended in CMM containing 20% Knock-out serum replacement and 10 µM ROCK inhibitor Y-27632 and sorted using a BD Fusion (BD Biosciences) into the same media. Cells were then collected by centrifugation at 200xg for 5 minutes.

### STARR-seq

#### Sequence Design

ATAC-seq peaks overlapping at least one genome-wide significant SNP (p < 5 × 10⁻⁸) from GWAS of atrial fibrillation, resting heart rate, or heart rate recovery post-exercise (20 s) were identified in SANCM clusters. To account for cluster-specific boundary variation, overlapping peaks were merged to generate a unified set of candidate regulatory regions. Peaks were further filtered to retain high-confidence regions using ArchR^88^ metrics (Peak_Max_Score > 200 and Peak_Min_Reproducibility > 2). From each retained peak, the central 500 bp was extracted from the hg38 reference genome. All SNPs within these regions were incorporated. For peaks containing a single SNP, two sequences were generated: one with the reference allele and one with the alternative allele. For peaks containing multiple SNPs, two composite sequences were designed: one harboring all alleles positively associated with the traits of interest, and the other with alleles negatively associated.

The nearest gene and its transcriptional orientation were used to define sequence strand. For genes on the positive strand, peaks upstream of or overlapping the transcription start site (TSS) were extracted from the negative strand, and those downstream from the positive strand. For genes on the negative strand, the opposite convention was applied. A total of 335 target regions (500 bp each; 167,500 bp in total) were selected, including 224 associated with atrial fibrillation, 107 with resting heart rate, and 32 with heart rate recovery post-exercise (20 s), with 28 regions overlapping between traits. To serve as negative controls, five sequences were randomly selected from non-repetitive genomic regions outside of ATAC-seq peaks, yielding 340 regions in total.

#### Library Preparation and Sequencing

STARR-seq was performed using the previously published protocol (Muerdter et al. Nat Methods. 2018) with the following details. Focused STARR-seq libraries were generated by synthesizing DNA fragments (Twist Bioscience) containing 500 bp regions centered around the scATAC-seq peak with overhangs for In-Fusion cloning (Takara) into AgeI and SalI digested hSTARR-seq_ORI vector (Addgene plasmid #99296). For screening, 6 × 10^6^ SANCMs were thawed, plated and allowed to recover as described above. Cells were then harvested using prewarmed Accumax (Sigma) for 20-30 minutes at 37°C. Digestion was quenched using CMM containing 20% Knock-out serum replacement (ThermoFisher) and 10 µM ROCK inhibitor Y-27632 and cells collected by centrifugation at 200xg for 5 minutes. Cells were resuspended at 8 × 10^6^ cells/ml in Amaxa P3 primary cell 4D-Nucleofector solution (Lonza) containing 4 µg of purified library per 1 × 10^6^ cells. Electroporation was carried out using an Amaxa 4D-Nucleofector X Unit (Lonza) by transferring 100 µl of above mix to each single Nucleocuvette and using the CA-137 Nucleofector program. Cells were then recovered and replated in CMM containing 20% Knock-out serum replacement (ThermoFisher) and 10 µM ROCK inhibitor Y-27632 for 24 hours. After 24 hours, media was changed to CMM for another 24 hours before cells were harvested for RNA isolation. Finished RNA-seq libraries were sequenced using an Illumina NextSeq2000 and PE300 reads.

#### STARR-seq data analysis

Paired-end sequencing (2 × 300 bp) was performed for one input library and five replicate STARR-seq libraries. Low-quality bases and adapter sequences were trimmed using *cutadapt* (v4.9; parameters: -q 20 -m 30) [https://doi.org/10.14806/ej.17.1.200]. Trimmed reads were aligned to the hg38 reference genome with *BWA-MEM* (v0.7.17) [https://arxiv.org/abs/1303.3997] using default settings. Reads with mapping quality <30 were removed with *samtools view* (v1.20) ^93^. Coverage was calculated using *bedtools genomecov*^90^with the -pc option, where coverage represents the number of fragments spanning each base. Fragment counts were summarized per target region and normalized to counts per million mapped fragments. For each replicate, log₂ ratios of STARR-seq signal versus input were calculated after adding a pseudocount of 1. Regions with an average log₂ ratio >0.585 across all five replicates were considered significantly enriched.

To identify SNPs with allelic differences in enhancer activity, two complementary approaches were used. First, allelic imbalance was assessed by counting reads supporting each allele using *bcftools mpileup* followed by variant calling with *bcftools call*^94^. Fisher’s exact test was applied to each replicate, and p-values were adjusted for multiple testing with the Benjamini–Hochberg method. Significant allelic imbalance was defined as FDR <0.05 and an absolute difference in reference allele frequency >10%. Second, for each SNP, read depth supporting each allele was normalized separately by counts per million mapped fragments, and log₂ fold changes (log₂FC) versus input were calculated for each replicate. Differences in log₂FC between alleles were tested using paired t-tests, with multiple testing correction by the Benjamini–Hochberg procedure. SNPs with FDR <0.05 were considered significant.

### Analysis of Chromatin Accessibility Allelic Imbalance

To identify variants exhibiting chromatin accessibility allelic imbalance (CAAI), we applied a modified approach based on a previously described method.^75^ First, we performed whole-genome sequencing (WGS) of the iPSC line at 30X coverage. Heterozygous single nucleotide variants (SNVs) were called from the WGS data. Sequencing reads from the scATAC-seq dataset were then separated by cell type. For each cell type, WASP (v.0.3.4)^95^ was used to remap scATAC reads to correct for allelic read-mapping bias. Reads aligned to heterozygous SNVs were counted using bcftools mpileup, followed by variant calling with bcftools call.^94^ SNVs with fewer than two reads from either allele or located outside of ATAC-seq peaks were excluded. The number of SNVs analyzed for allelic imbalance was 44,316 in SAN tail cells, 33,043 in SAN head cells, 38,671 in transitional cells, and 38,204 in sinus venosus cells. Allelic imbalance was assessed using a two-sided binomial test with multiple testing correction applied via the Benjamini–Hochberg method and a false discovery rate (FDR) cutoff of 0.1.

To investigate transcription factor motifs at sites with CAAI, we used FIMO (v.5.5.8)^96^ to scan the 20 bp region surrounding each variant against the motifs in the JASPAR 2022 CORE non-redundant vertebrates database^97^. Both the reference and alternate sequences were queried, retaining motif matches with P-values below 10^−3^ for either allele. Only matches with log10|p-value_ref_/p-value_alt_| > 1 were analyzed. Enrichment of motifs impacted by CAAI variants was evaluated by comparison with a set of read count-matched variants lacking CAAI (tenfold larger), using a one-sided Fisher’s exact test. P-values were corrected for multiple hypothesis testing using the Benjamini–Hochberg procedure. To investigate the effect of transcription factor motifs on the directionality of chromatin accessibility, we examined each motif predicted to be altered by a CAAI variant. Specifically, we determined the proportion of motif occurrences where the allele exhibiting the stronger motif match (indicated by the lower FIMO P-value) was also associated with increased chromatin accessibility.

### Measurement of SANCM Beat Rate in the Presence of Carbachol

SANCMs were recovered and plated in 35 mm dishes containing 14 mm glass microwells (MatTek). On the day of transfection media was changed to CMM with 10% FBS. Transfections were carried out using Viafect transfection reagent (Promega) at a ratio of 6:1 (µl Viafect: µg DNA). Transfection complexes were formed in Opti-MEM serum free media (ThermoFisher) according to the manufacturer’s instructions using 1.6 ug of pcDNA3.1+ or pcDNADNA3.1+ containing the coding sequence for GNB4 per dish. 24 hours post-transfection, media was changed to CMM for an additional 24 hours before beat rate was assessed using IonOptix video-based edge detection system. Cells were maintained at 37°C in Tyrode’s solution containing: 140 mM NaCl, 5.4 mM KCl, 1 mM MgCl₂, 1.8 mM CaCl₂, 10 mM glucose, and 10 mM HEPES (pH 7.4). Prior to recordings, SANCMs were equilibrated for 10–15 minutes before contractile measurements were acquired using the IonOptix MyoCam camera mounted on an inverted microscope. Spontaneously beating SANCMs were visualized under phase-contrast illumination and cellular edge displacement was recorded at high frame rates. Baseline recordings were obtained prior to the addition of indicated concentrations of Carbachol (Sigma). Between doses, SANCMs were washed 3x with 37°C Tyrode’s then equilibrated for 5 minutes before measurements were continued. Data acquisition was performed using IonWizard software and analysis was performed using CytoSolver software.

## Acknowledgements

The authors wish to thank members of the Vedantham Lab for helpful discussions.

---

**Extended Data Figure 1:**
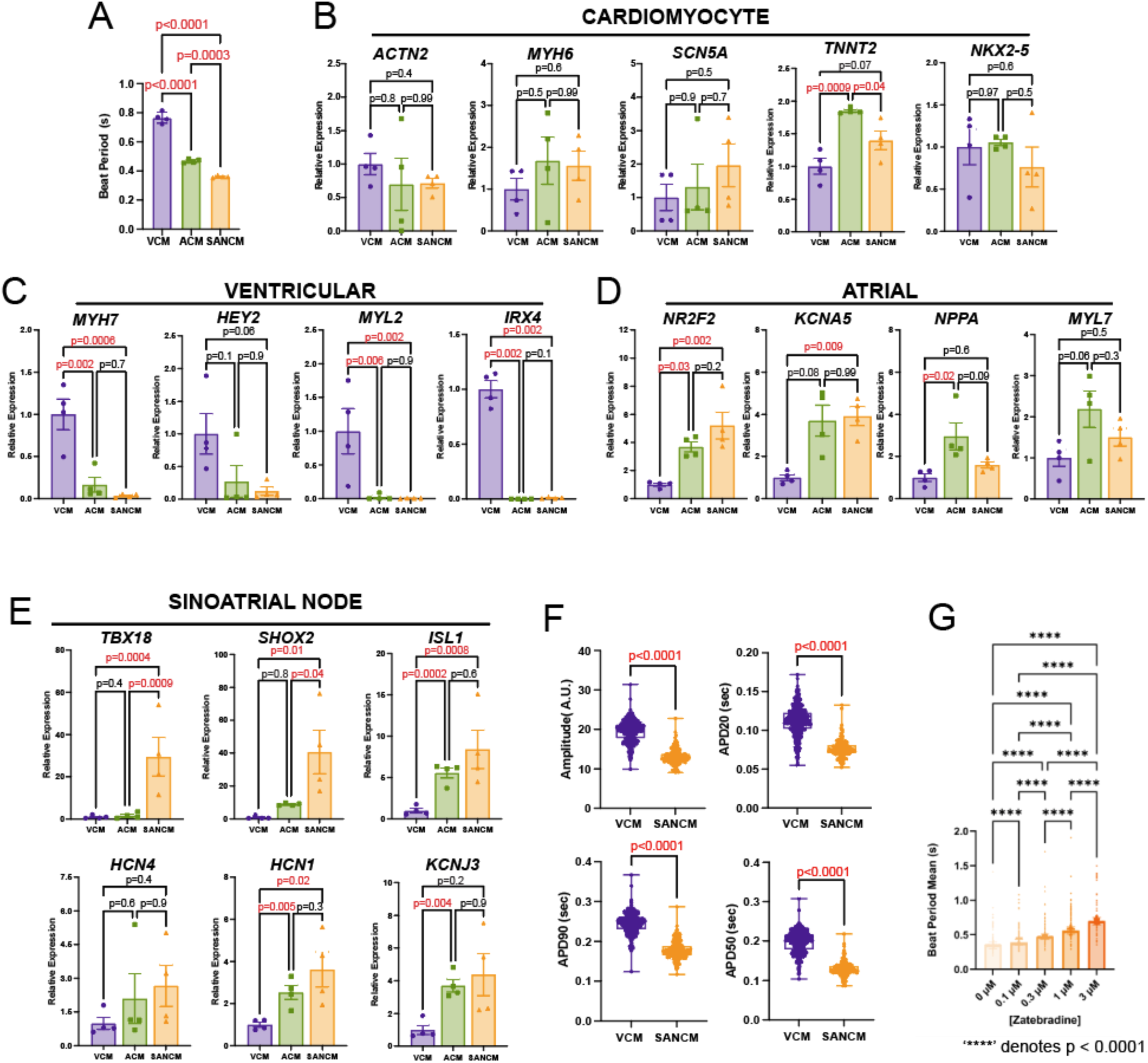
Characterization of hiPSC derived Cardiomyocyte-like subtypes. (A) Beat period as measured by MEA in 4 separate monolayer differentiations of hIPSC-derived ventricular (VCM), atrial (ACM) and sinoatrial node (SANCM) cardiomyocytes. (B-E) quantitative PCR for (B) pan-cardiac, (C) ventricular, (D) atrial genes, and (E) pacemaker associated genes in VCMs, ACMs, and SANCMs. For quantitative PCR, the ΔΔCT method was used with all values normalized to the average of the VCM differentiations. Statistical tests were performed with one-way ANOVA and post-hoc adjustments of P value for multiple tests. Significant P values are shown in red. (F) Optical action potentials parameters measured using FluoVolt including amplitude (top, left panel), action potential duration at 20, 50, and 90 percent repolarization (APD 20, APD 50, and APD90, respectively for groups of single VCM and SANCM cells. Comparison was made with a t-test and significant P values are shown in red. (G) Changes in beating period of SANCMs in response to escalating doses of zatebradine, a specific blocker of Hcn4 channels. One-way ANOVA with post-hoc adjustments of P values for individual comparisons was used to compare groups. All individual comparisons were significant.

**Extended Data Figure 2:**
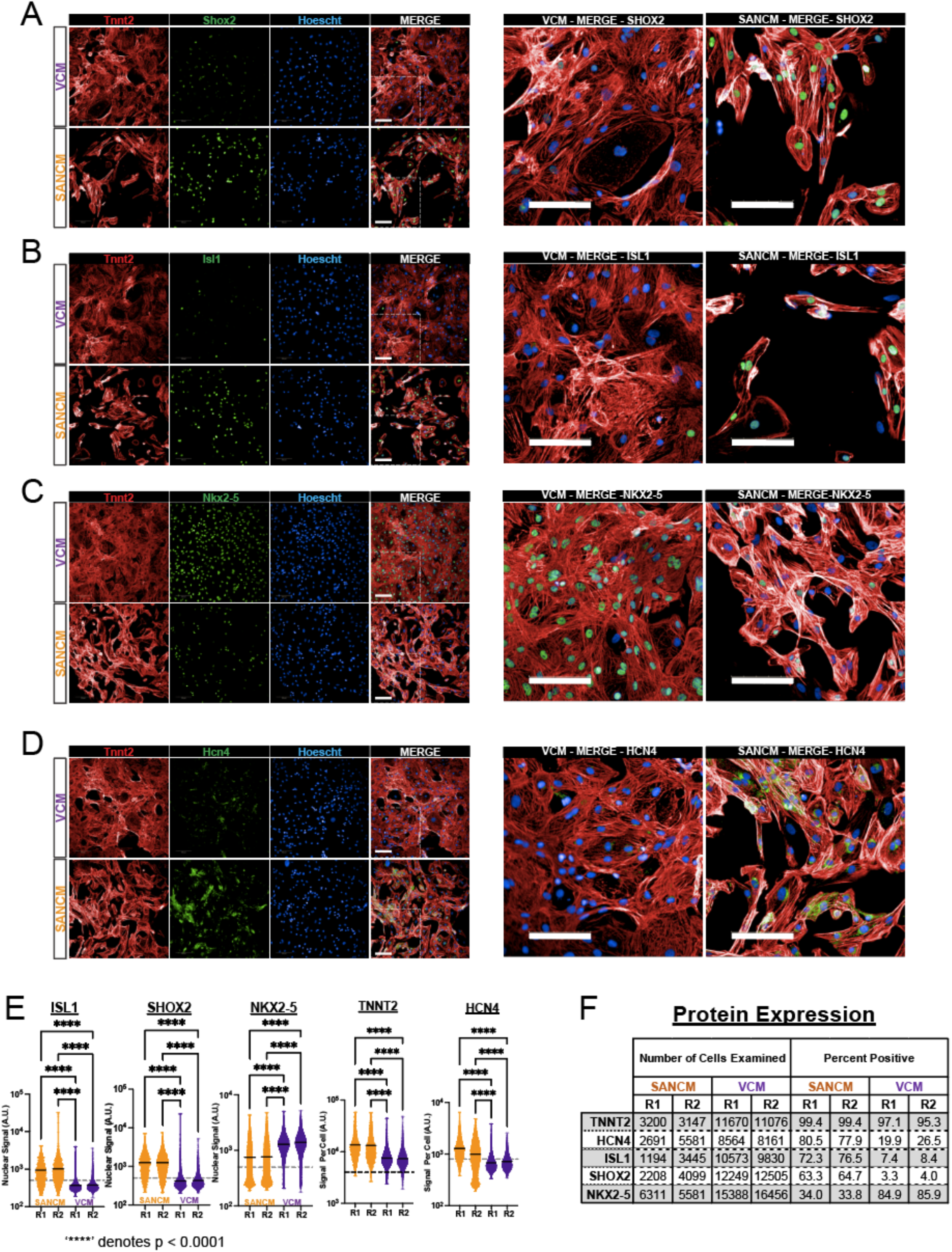
Expression of Lineage-Defining Proteins in hiPSC deviced Cardiomyocyte-like Subtypes. Immunocytochemistry for either (A) SHOX2, (B) ISL1, (C) NKX2-5, or (D) HCN4 (green) with TNNT2 (red) and hoescht (blue) in hiPSC-derived sinoatrial node-like cardiomyocytes (SANCMs) compared to ventricular-like cardiomyocytes (VCMs). VCMs are displayed in the top rows and SANCMs in the bottom rows, with areas of detail from the merged images expanded to the right. Scalebars = 100 μm. (E) Quantification of signal intensity from immunocytochemistry images for SANCMs and VCMs from randomly selected high-power fields from 2 separate differentiations. ‘R1’ denotes replicate 1 and ‘R2” denotes replicate 2. The dashed line indicates empirically defined thresholds used for scoring cells as either positive or negative for the indicated gene. Statistical significance was assess with ANOVA, followed by post-hoc pairwise comparisons (‘****’ denotes P < 0.0001) (F) Table summarizing the quantification of immunocytochemistry images, with the number of cells examined and the percentage with signal intensity for the indicated protein that fell above the empirically defined threshold shown in (E).

**Extended Data Figure 3.**
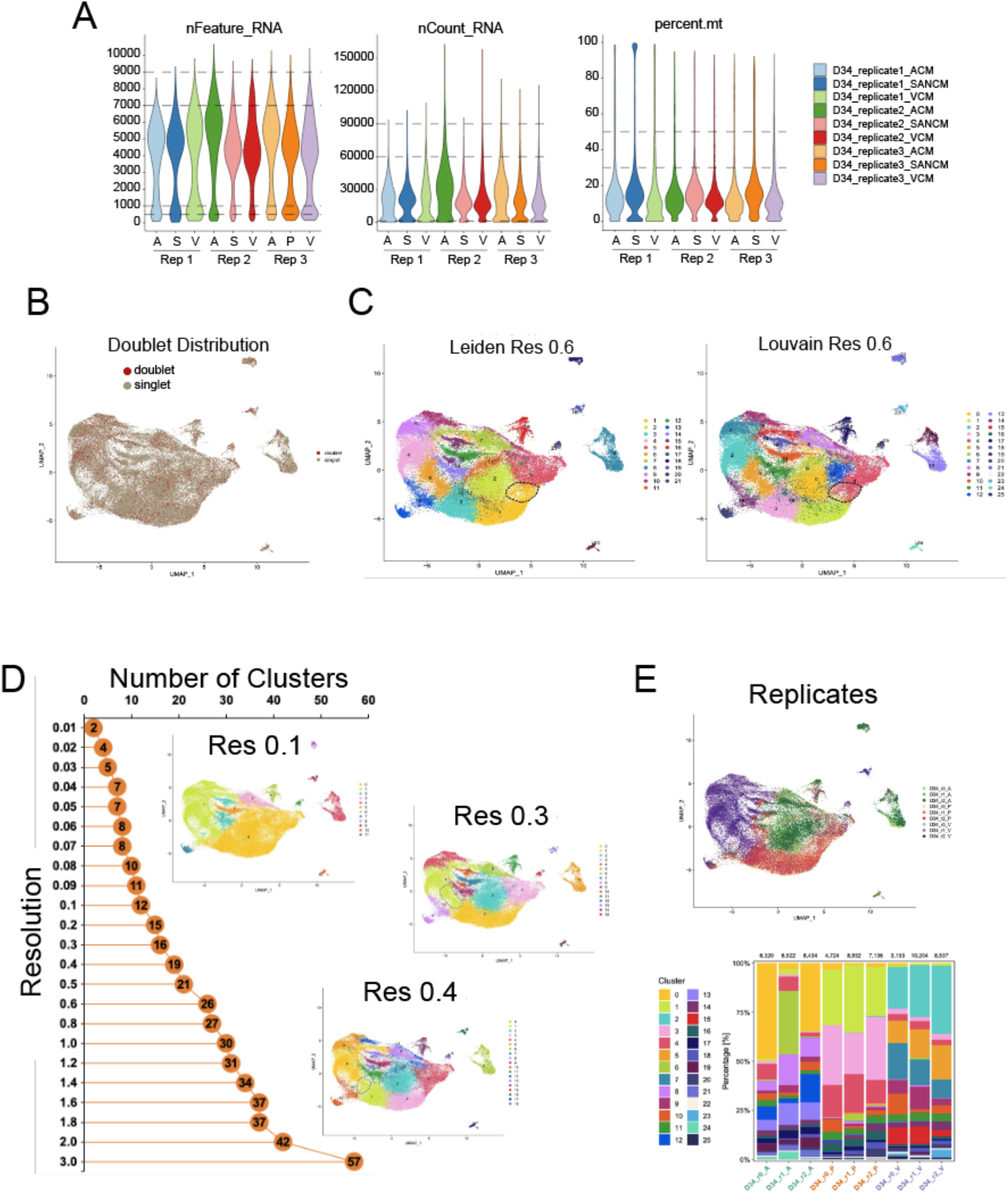
RNA Sequencing and Clustering of HiPSC-Derived Cardiomyocytes. (A) Violin plots that display (left) the number of genes with mapped reads per cell, (center) the total number of mappable reads per cell, and (left) the percentage of reads that align to the mitochondrial genome across all samples. (B) Uniform Manifold Approximation and Projection (UMAP) showing that the distribution of doublets (red) is uniform. (C) UMAPs comparing Leiden clustering and Louvain clustering at the chosen resolution of 0.6. (D) The number of clusters as a function of clustering resolution with UMAPs displayed corresponding to resolutions of 0.1, 0.3, and 0.4. (E) UMAP partitioned by replicate (top) with quantification of cluster assignments by replicate below.

**Extended Data Figure 4.**
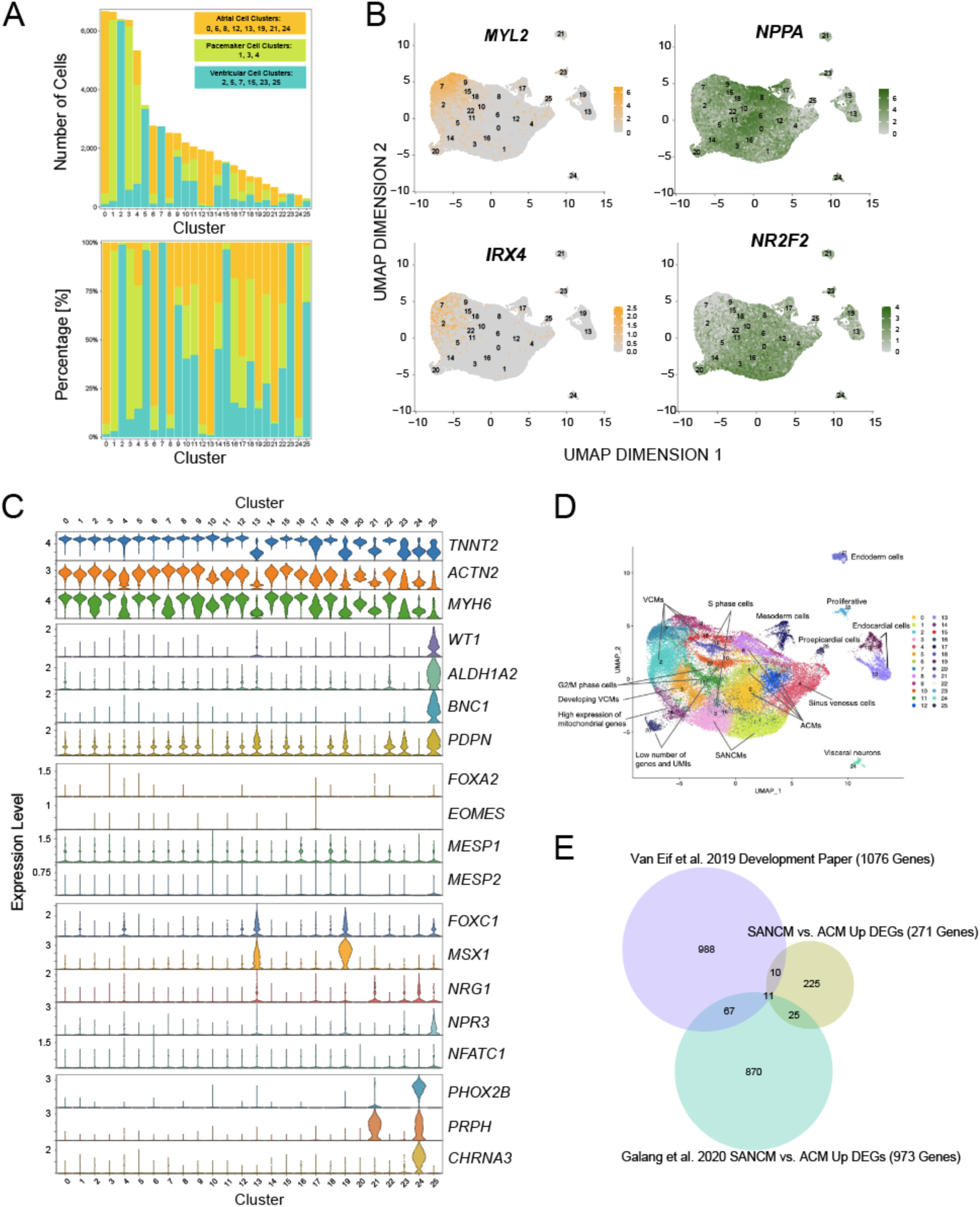
Cluster Annotation for hiPSC-Derived Cardiomyocytes. (A) *Top panel*, number of cells falling into each cluster, colored by cell type of origin. *Bottom panel*, Cluster composition expressed as percentage of cells in each cluster from each cell type of origin. (B) Uniform manifold approximation and projection (UMAP) for combined Ventricular (VCM), atrial (ACM) and sinoatrial (SANCM) cardiomyocytes demonstrating expression of 2 ventricular marker genes (MYL2 and IRX4) predominantly in VCMs, and atrial marker gene NPPA most strongly in ACMs and atrial/sinoatrial gene NR2F2 predominantly in ACMs and SANCMs. (C) Violin plots for gene expression by cluster for makers of cardiomyocytes (*TNNT2, ACTN2, MYH6*), proepicardium (*WT1, ALDH1A2, BNC1, PDPN*), endocardium (*FOXC1, MSX1, NRG1, NPR3*, *NFATC1*), mesendoderm (*FOXA2, EOMES, MESP1*, *MESP2*), and visceral neurons (*PHOX2B, PRPH, CHRNA3*) (D) UMAP with labelled by cluster annotations based on marker gene expression. (E) Venn Diagram demonstrating overlap among differentially expressed genes for 3 datasets comparing sinoatrial cardiomyocytes to atrial cardiomyocytes: SANCMs versus ACMs from the present study, sorted *Hcn4-GFP^+^* neonatal SAN cells versus *Myh6-mCherry^+^* atrial cardiomyocytes from Galang et al (2020) and sorted Tbx3+ cells versus atrial cells from Van Eif et al. (2019).

**Extended Data Figure 5.**
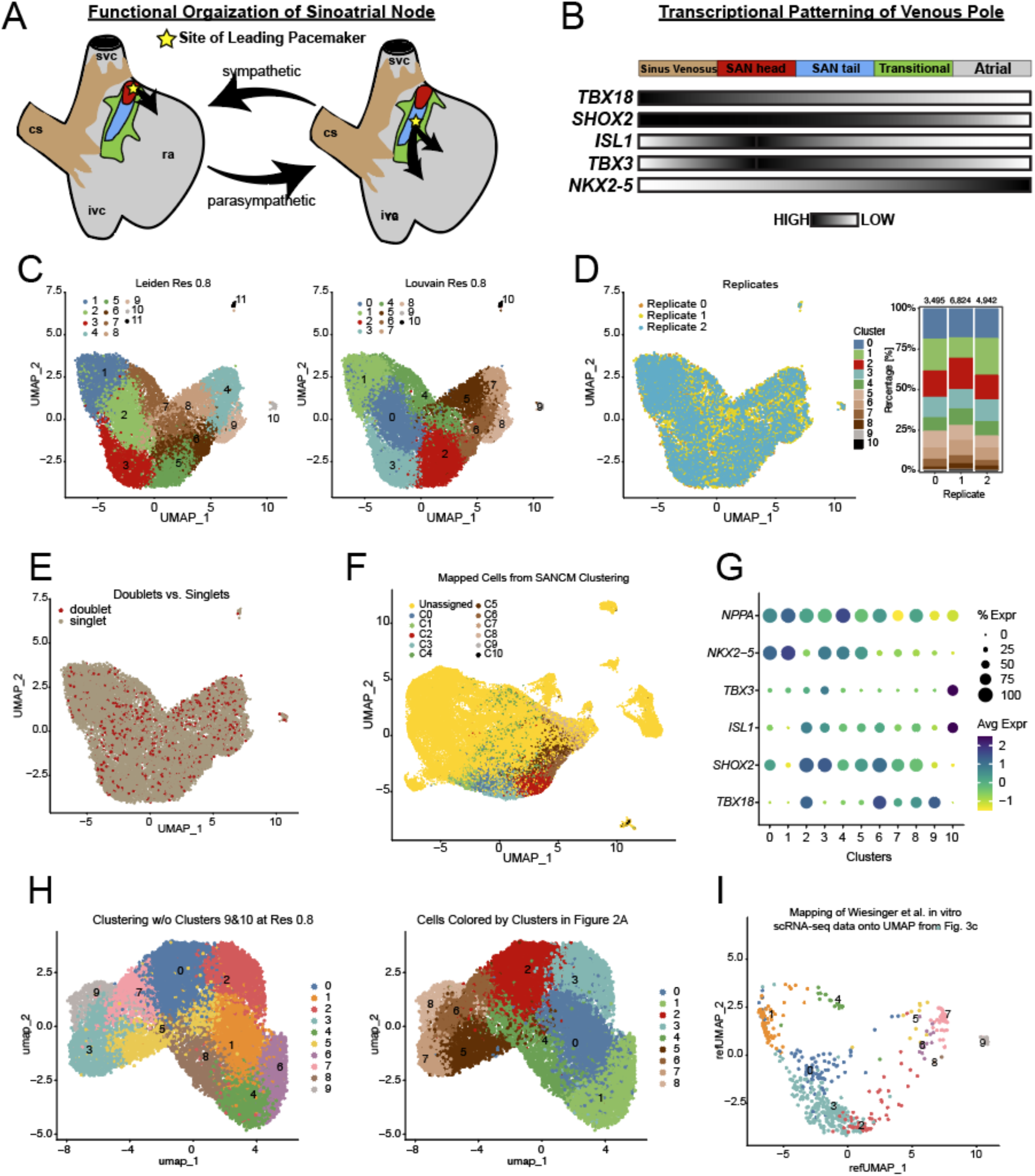
Sub-Clustering HiPSC-Derived SAN-like Cardiomyocytes. (A) Illustration showing the anatomical and functional organization of the Sinoatrial Node (SAN). When sympathetic tone is high, the cranial region of the SAN head (red) is the site of the leading pacemaker; when parasympathetic tone predominates, the SAN tail (blue) is the site of the leading pacemaker. Transitional myocardium (green) and sinus venous myocardium (brown) are also shown. Abbreviations: svc; superior vena cava, cs; coronary sinus, ivc; inferior vena cava, ra; right atrium. (B) Diagram depicting the gradient expression of key transcription factors in the patterning of the myocardia subtypes at the cardiac venous pole. Darker color indicates a higher expression level. (C) Uniform manifold approximation and projection (UMAP) with Leiden (left) versus Louvain (right) clustering at resolution 0.8 shows minor differences. (D) UMAP with annotation by replicate (left) and a plot (right) showing the cluster composition by replicate. There were no significant differences among the replicates. (E) UMAP with doublet distribution in the SANCMs (F) The combined UMAP for ACMs, VCMs, and SANCMs, colored by SANCM subclusters demonstrates that SANCM subclusters including SAN head, SAN tail, transitional and sinus venosus cells fall into largely separate regions. (G) Bubble plot showing expression of key cardiac transcription factors and marker genes in each SANCM subcluster (H) Re-clustering of SANCMs after exclusion of non-cardiomyocyte clusters is shown in the UMAP at left. The UMAP in the right panel shows the same clustering but colored according to the original clustering, demonstrating that including non-myocytes in the original clustering did not have a substantial impact on clustering of cardiomyocytes. (I) Projection of cells from Weisinger et al (2022) into the SANCM UMAP demonstrates that most cells map into different pacemaker subclusters with SAN tail most strongly represented.

**Extended Data Figure 6:**
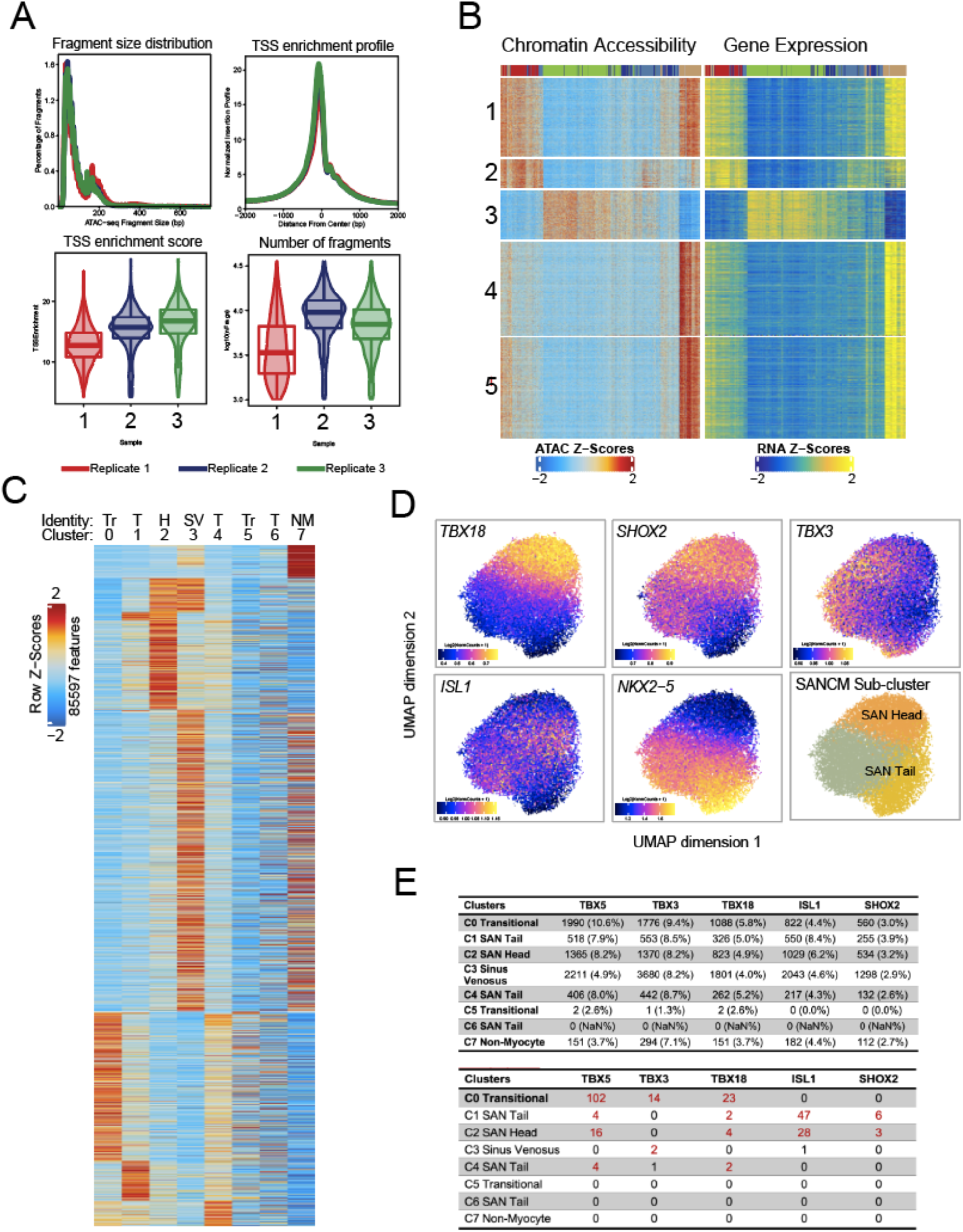
Single cell ATAC-seq Analysis of SAN-like Cardiomyocytes. (A) Summary of scATAC-seq quality control metrics for each SANCM replicate including fragment sizes and transcription start site (TSS) enrichment profile. (B) Heatmap summary of ArchR peak-to-gene linkages between scATAC and scRNA-seq data of SANCMs (correlation≥0.45). (C) Heatmap of marker peaks for each cluster shown in Fig. 3a demonstrates distinct profiles for each cluster. Abbreviations: Tr, transitional; T, SAN tail; H, SAN head; SV, sinus venosus; NM, non-myocyte. (D) Feature plots showing predicted expression of SANCM canonical genes in sub-clustering analysis of SAN Head (C2) and SAN Tail (C1, C4, C6) clusters from Fig. 3a. Note that NKX2-5 and TBX-18 prediction expression are complementary in the UMAP as expected from the corresponding RNA-seq data shown in Fig. 2d (E) Motif enrichment analysis for SAN transcription factors for marker peaks in each cluster. The top table shows the number of peaks (percentage in parenthesis) that contain motifs for each transcription factor indicated in the top row. The bottom table shows the -log (10) adjusted P value for each motif for marker peaks in each cluster. Note that ISL1 motifs are enriched in SAN head and SAN tail. Adjusted P values < 0.05 are deemed significant and are shown in red.

**Extended Data Figure 7:**
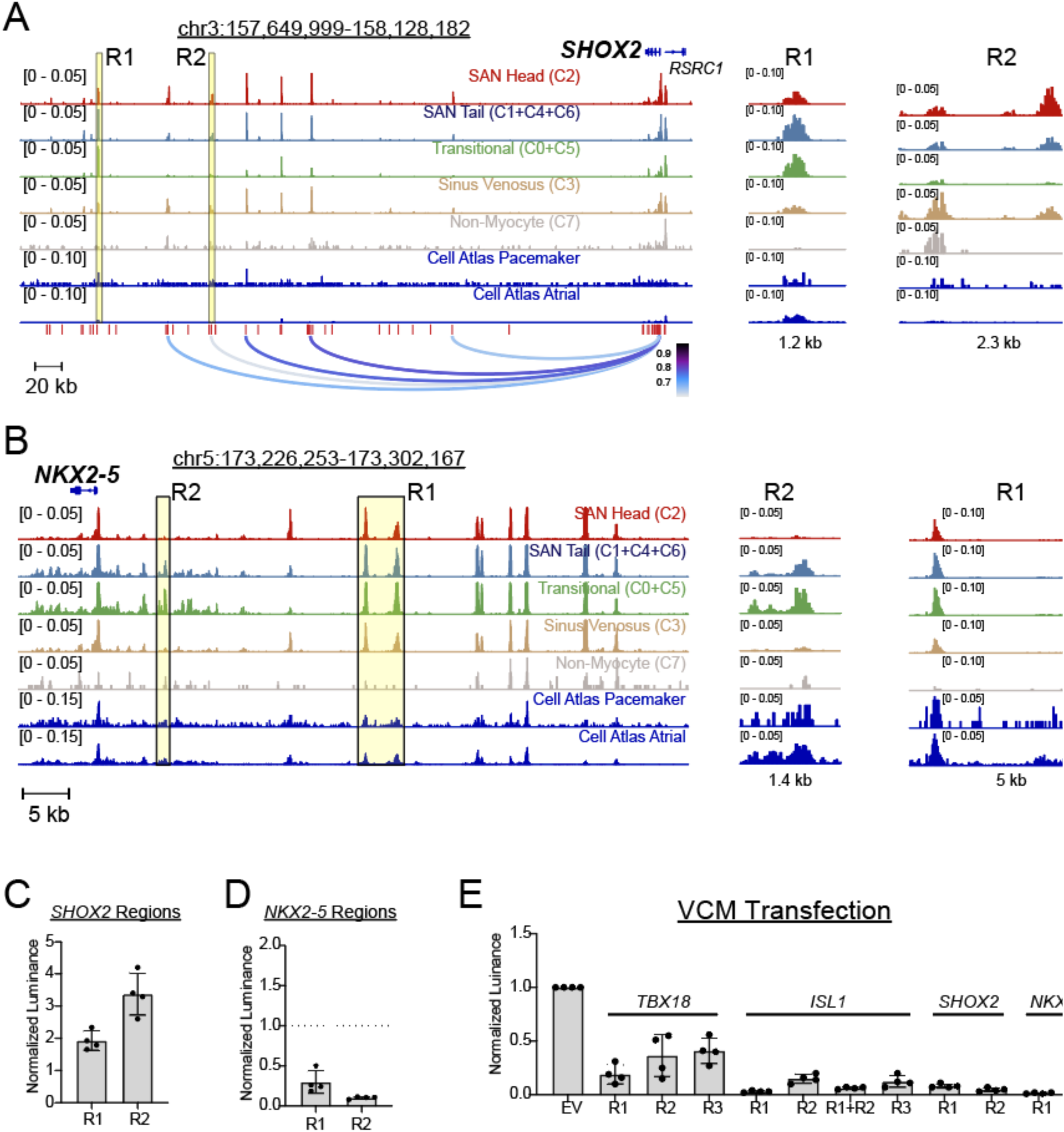
Regulatory elements at SHOX2 and NKX2-5 loci. (A) Chromatin accessibility tracks at the *SHOX2* locus from SANCM scATAC-seq re-bulked by cluster annotations, with re-bulked scATAC-seq tracks from Heart Cell Atlas pacemaker cells and atrial cardiomyocytes below. Peak-to-gene (P2G) links defined by applying ArchR to SANCM scATAC-seq and scRNA-seq are shown at the bottom with the color scale representing correlation strength. Two accessible chromatin regions (R1 and R2) are highlighted in yellow and shown at higher magnification in the panels to the right. Peak heights were normalized and scaled by the ReadsInTSS parameter, the default normalization method in ArchR (B) Browser tracks from the *NKX2-5* locus analogous to those displayed in panel A that highlight two regulatory elements with differential accessibility among different clusters were cloned for functional analysis. (C) Luciferase/Renilla reporter activity for 2 plasmid constructs generated for *SHOX2-R1* and *SHOX2*-*R2* are shown normalized by activity of the empty luciferase vector in transfected SANCMs (n = 4 biological replicates). *P* values are shown for comparison to a value of 1 corresponding to empty vector. (D) Luciferase/Renilla reporter activity for 2 *NKX2-5-R1* and *NKX2-5-R2* and shown normalized by activity of the empty luciferase vector in transfected SANCMs (n = 4 biological replicates). *P* values are shown for comparison to a value of 1 corresponding to empty vector. The *NKX2-5* elements both result in repression of basal promoter activity in SANCMs. (E) Luciferase/Renilla reporter activity for empty vector and regulatory elements at *TBX18*, *ISL1*, *SHOX2*, and *NKX2-5* show repression of basal promoter activity when transfected into VCMs (n = 4 biological replicates).

**Extended Data Figure 8.**
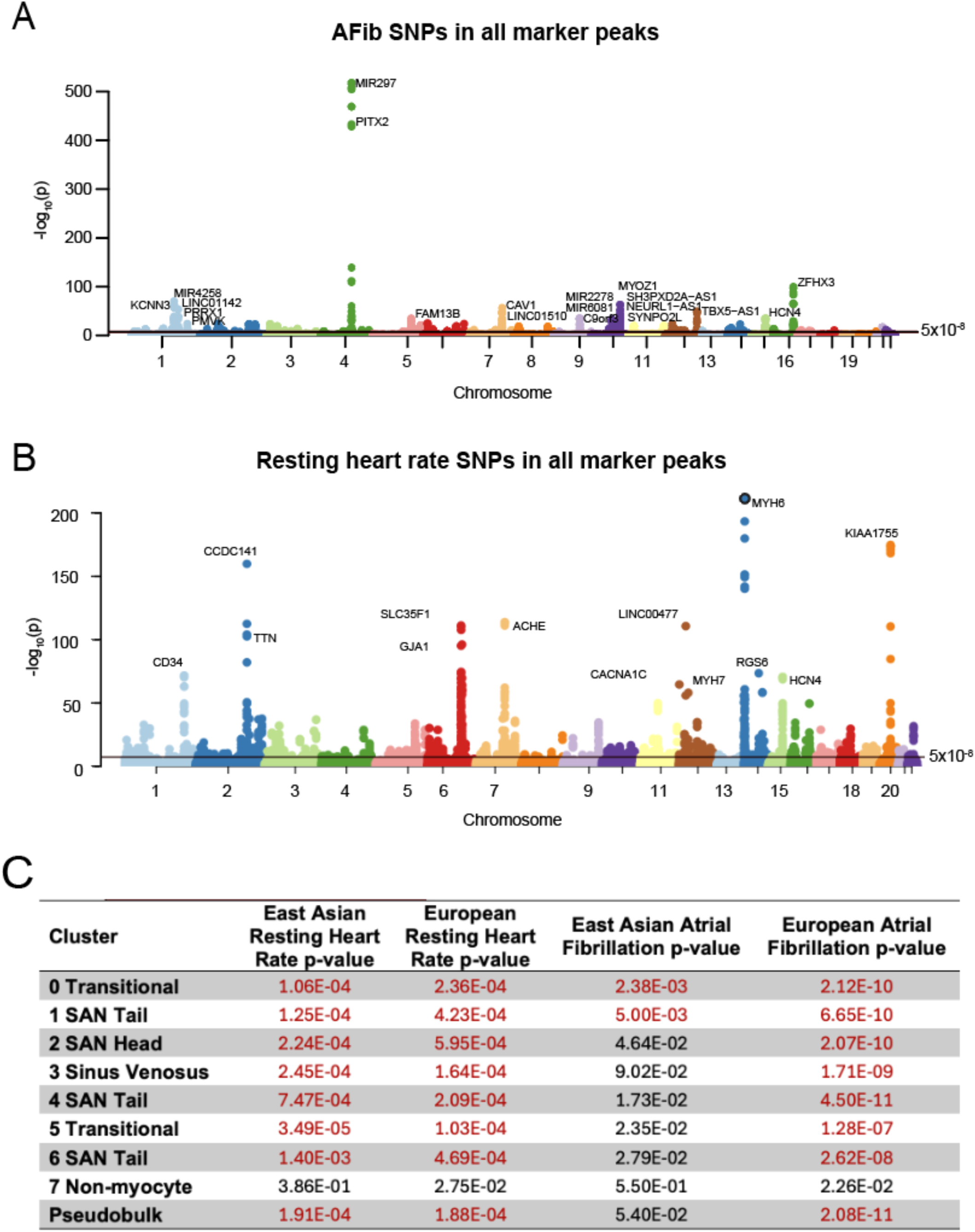
Trait Associations for Single Cell ATAC-seq Based Clusters. Manhattan plot showing significant genomic associations with atrial fibrillation (A) and resting heart rate (B) that overlap with scATAC-seq maker peaks (as opposed to all peaks) (C) Adjusted P values for association with atrial fibrillation and resting heart rate for single nucleotide polymorphisms within open chromatin regions in scATAC-seq cluster using datasets from European individuals and East Asian individuals. Despite the reduced power for East Asians due to lower sample size (4,150 cases in East Asians versus 60,620 in Europeans) findings are broadly concordant among the 2 ancestries.

**Extended Data Figure 9.**
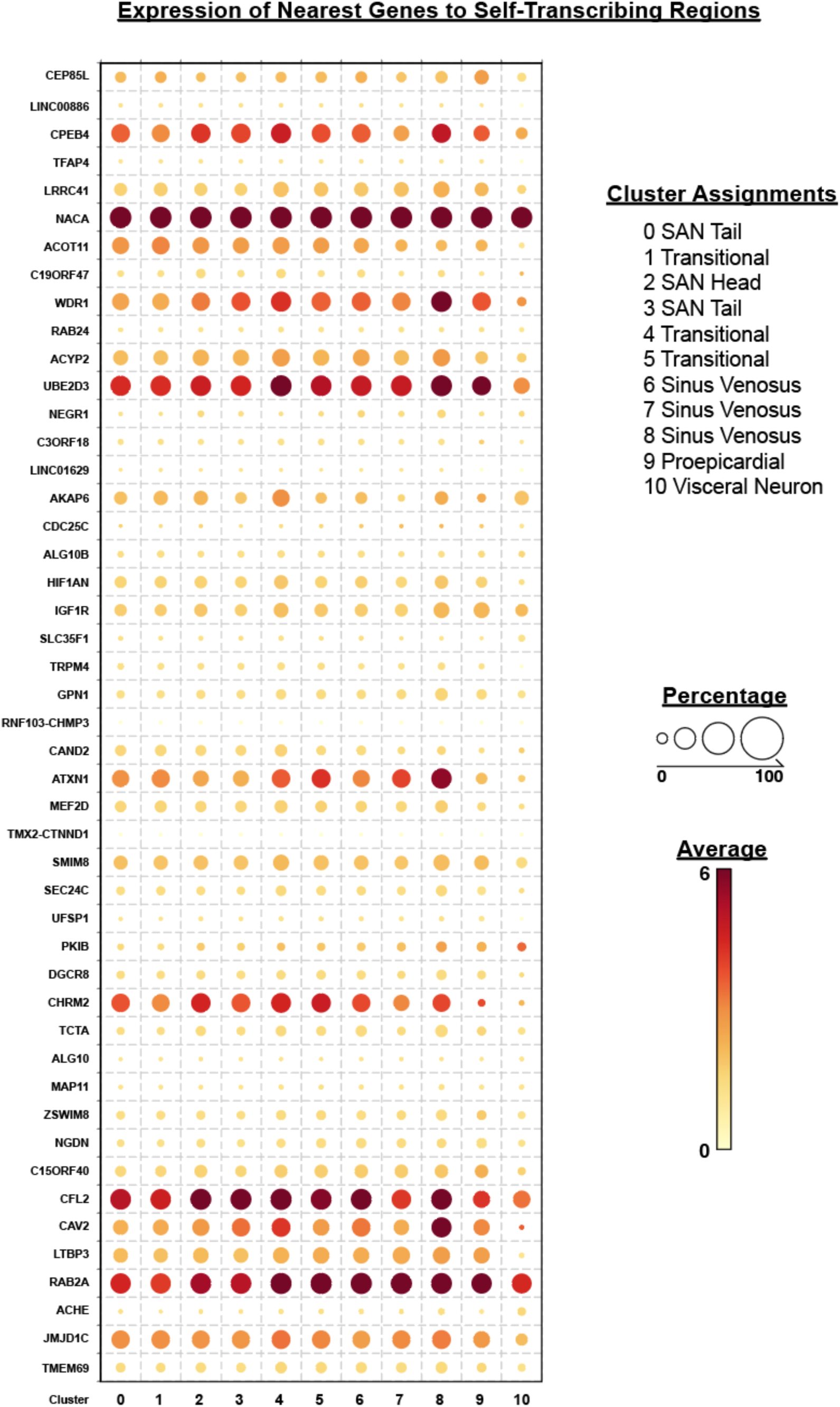
Expression of Nearest Genes to Self-Transcribing Regions. Bubble plot showing expression level by color scale and percentage of cells positive for expression by bubble size in each scRNA-seq cluster for each of the nearest genes to the top 48 scATAC-peaks with greater than 1.5-fold change in mRNA/DNA in the self-transcribing active regulatory region sequencing (STARR-seq) experiment shown in Figure 6. Note that while *ATXN1*, an AF-associated gene, demonstrates expression in all cardiomyocyte clusters, it’s expression is highest in sinus venosus where AP-1 is an upstream regulator based on motif enrichment in scATAC-seq peaks.

